# Disentangling multivariate relationships between cognition, language and social traits: structures of G, E, and r_GE_

**DOI:** 10.1101/2025.07.26.666154

**Authors:** Fenja Schlag, Lucía de Hoyos, Ellen Verhoef, Alexander Klassmann, Simone van den Bedem, Simon E. Fisher, Brad Verhulst, Beate St Pourcain

## Abstract

**Background:** Cognitive, language, and social abilities are complex, heritable and intertwined traits shaping children’s development and later mental health. To better understand cross-trait interrelationships, we model here the structures of shared genomic and shared non-genomic/residual (i.e. broadly environmental) influences, and their correlation (r_GE_), investigating cognitive, language, and social behavioural/communication measures.

**Methods:** Data were obtained for unrelated children (8-13 years) from two population-based cohorts: the UK Avon Longitudinal Study of Parents and Children (ALSPAC, N≤6,543) and the US Adolescent Brain Cognitive Development℠ (ABCD) Study (N≤4,412), and analyses were carried out implementing an extended data-driven genetic-relationship-matrix structural equation modelling (GRM-SEM) approach.

**Results:** In ALSPAC, we identified two independent phenotypic domains, each captured by a structurally matching pair consisting of a genomic (A) and a non-genomic/residual (E) factor. The first domain reflected cognitive/language difficulties, with the largest genomic and residual factor loadings (λ_A_ and λ_E_, respectively) for verbal IQ (λ_A_=0.73(SE=0.05); λ_E_=0.57(SE=0.07)). The second domain captured social difficulties, with the largest λ_A_ and λ_E_ for social communication measures (λ_A_=0.39(SE=0.10); λ_E_=0.82(SE=0.10)). We identified trait-specific r_GE_ between pairs of A and E factors with different directions of effect (cognition/language r_GE_=0.89(SE=0.18), social r_GE_=-0.62(SE=0.17)). r_GE_ patterns were linked to increased measurable A and E contributions for cognition/language difficulties, but decreased contributions for social problems. Analyses in ABCD confirmed the two domains for E and phenotypic structures, although genomic contributions were low.

**Conclusions:** In childhood, cognitive/language abilities versus social abilities are influenced by distinct genomic and/or environmental factors, potentially interlinked through trait-specific r_GE_, suggesting differences in developmental processes.

## Introduction

Cognitive, language, and social abilities are complex, interrelated (1,2) traits that crucially shape children’s development, educational outcomes and mental health (1,3–6). Childhood cognitive abilities support the acquisition of knowledge, information processing and reasoning, and are linked to both language development (7) and social cognition (8). Productive and receptive language abilities enable verbal interactions with others, facilitating the exchange of meaning and thought (9) and rely on a broad spectrum of social skills (10). Social abilities may encompass intentional social actions (11) but also social communication, including verbal or non-verbal interactions, and the interpretation of language within a social context (pragmatics)(12). Developmental interrelationships between these domains may involve education-related cascading effects, especially from mid-childhood onwards (13,14) affecting wellbeing across the life course(6,15).

Variation in cognitive, language, and social abilities can partially be attributed to genetic influences and partially to environmental (non-genetic) influences although the structure of variance patterns is little characterised. Genome-wide studies tagging variation in common single-nucleotide polymorphisms (SNPs), reported SNP-h^2^ estimates of 17% for cognitive abilities (16), 8-26% for language-related measures (17,18), and 3-47% for social traits (19–21). The remaining variation is typically attributed to environmental influences. These may, for example, act through educational and socio-economic resources that are nested within families and directly affect an individual’s development (22,23). Additionally, environmental differences can manifest within neighbourhoods and/or school districts (22) or through peers, life events, and social support (24).

Genetic and environmental influences may correlate due to passive (e.g., parental genetic factors creating their child’s environment), active (e.g., a genetic motivation to seek a particular environment) or evocative (e.g., genetically motivated behaviours eliciting reactions from others) mechanisms (gene-environment correlations, r_GE,_) (25). Recent genomic studies investigating educational attainment- and socioeconomic status-related traits primarily found evidence for passive r_GE_ through comparisons of polygenic load across family members (26–28), including extended family structures (29). These findings demonstrate that r_GE_ may arise within families and through dynastic social processes. However, studies of r_GE_ in unrelated individuals that do not rely on predefined nesting (family-based) structures and, consequently, may capture all types of r_GE_ are still outstanding.

To study inter-trait relationships across cognition, language, and social skills in unrelated children and young adolescents, we model here shared genomic and non-genomic (residual) structures and assess evidence for their correlation (r_GE_) using genetic-relationship-matrix (GRM) structural equation modelling (GRM-SEM) (19,30). To this end, we investigate data from two independent population-based samples: the UK Avon Longitudinal Study of Parents and Children (ALSPAC, N≤6,543)(31) and the US Adolescent Brain Cognitive Development^℠^ Study (ABCD Study®, N≤4,412) (32) (Figure 1). While prominent multivariate genomic modelling techniques rely on summary statistics derived from genome-wide association studies (33), GRM-SEM utilises direct individual-level genotype and phenotype information to model genomic and non-genomic co-variance structures (30). Extending our previous GRM-SEM framework (30), we study here correlation between genomic and non-genomic factors (r_GE_), without the need of predefined family-based structures.

**Figure 1:**
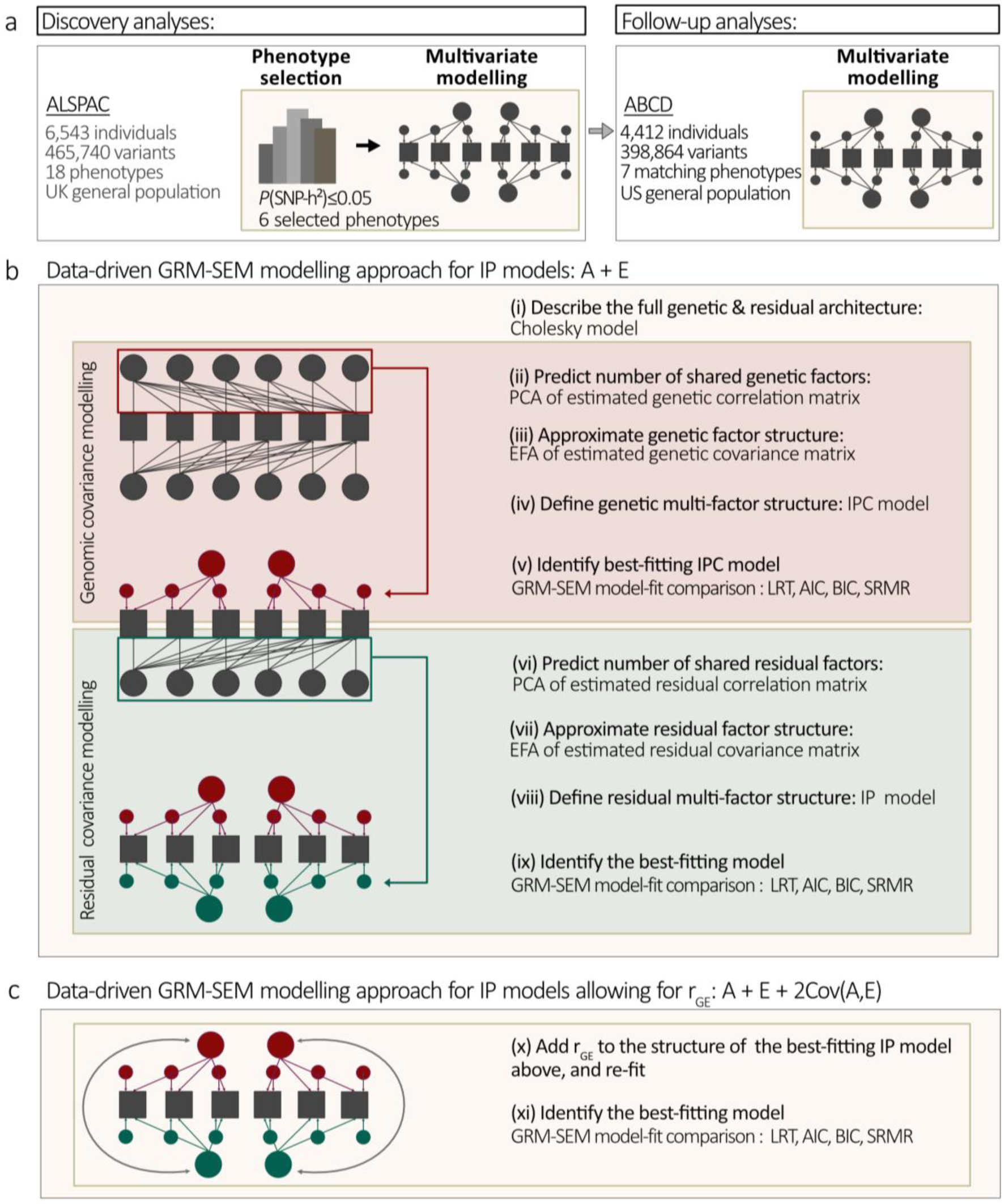
Study outline and modelling approach. **(a)** Study outline. Discovery analyses were performed in ALSPAC and followed up in the ABCD Study^®^. (b) Data-driven stepwise multivariate modelling approach of genomic and non-genomic structures. Note that for the ABCD Study^®^, due to low SNP-h², multivariate modelling was limited to the non-genomic structure, implementing a CIP model only. (c) Estimation of gene-environment correlations (r_GE_) extending the data-driven approach in (b). Given the similarity of genomic and non-genomic structures (ALSPAC only), r_GE_ parameters can be added to the identified IP from (b), to provide starting values for re-fitting the model. Note that there is evidence for r_GE_, if the IP model allowing for r_GE_ fits the data better than the IP model assuming independence of A and E (b), and the r_GE_ estimates are different from zero. All models are described in the Methods. Cholesky, IPC and CIP models are also schematically illustrated in Supplementary Figure 1, and the IP model with and without r_GE_ in Figure 2. ABCD - Adolescent Brain Cognitive Development Study (ABCD Study®); ALSPAC - Avon Longitudinal Study of Parents and Children; CIP - Hybrid Cholesky/independent pathway; EFA – Exploratory factor analysis; GRM-SEM - Genetic-Relationship-Matrix Structural Equation Modelling; IP - Independent pathway; IPC - Hybrid independent pathway/Cholesky; PCA – Principal component analysis; r_GE_ - Gene-environment correlation; SNP-h² - SNP heritability

## Methods

### Cohort information

<u>ALSPAC</u> is a UK population-based longitudinal pregnancy-ascertained birth cohort with birth dates between 1991 and 1992 (34,35) (<u>data dictionary and variable search tool</u>). Ethical approval for the study was obtained from the ALSPAC Ethics and Law Committee and the Local Research Ethics Committees. Consent for biological samples has been collected in accordance with the Human Tissue Act (2004). Informed consent for the use of data collected via questionnaires and clinics was obtained from participants following recommendations of the ALSPAC Ethics and Law Committee at the time (Supplementary Methods).

<u>The ABCD Study</u>**^®^** is a US population-based longitudinal study of brain development and child health (32). Starting at the ages of 9-10 years, participants were followed for 10 years at 21 data acquisition sites across the US (N=11,877, Data Release 4.0 (36)). The Institutional Review Board (IRB) at the University of California, San Diego, approved all aspects of the ABCD Study**^®^**(37). Parents or guardians provided written consent, while children provided written assent (38).

### Phenotype information

<u>ALSPAC:</u> A total of 18 mid-childhood/early adolescent measures (8-13 years) was included in the study (Supplementary Methods, Supplementary Table 1). <u>Cognition:</u> Children’s verbal and performance intelligence quotient scores (VIQ and PIQ respectively) were assessed with the Wechsler Intelligence Scale for Children (WISC) III (39) at age 9. <u>Language:</u> Children’s listening comprehension (LGC) was measured using a subtest of the Wechsler Objective Language Dimensions(40) at age 9. <u>Social</u> <u>traits</u>: Social reciprocity and verbal/nonverbal communication (social communication difficulties, SCD)(41) at 8 and 11 years were obtained with the Social and Communication Disorder Checklist using parent report. In addition, children’s social communication at 10 years was assessed with parent-reported pragmatic composite scores (PRC) of the Children’s Communication Checklist (CCC) (42). Social behaviour, as captured with the prosocial behaviour (PB) and peer problems (PP) subscales of the Strengths and Difficulties Questionnaire (SDQ)(43), was measured at the age of 7, 10, 12, and 13 years based on parent report, and at the age of 8 and 11 years based on teacher report.

<u>ABCD:</u> Using phenotype release 4, a total of 7 mid-childhood/early adolescent measures were studied (Supplementary Methods, Supplementary Table 2). <u>Cognition:</u> Crystallised (CC) and fluid (FC) cognition composite scores were evaluated at 10 years using the NIH Toolbox (44). Matrix reasoning (MR), as a measure of non-verbal reasoning, was assessed at the age of 10 years using the WISC-V (45). <u>Social traits:</u> Social behavior was measured with the social problems (SP) subscale of the Child Behaviour Check List (46) at 10, 11, 12, and 13 years based on parent reports.

### Phenotype transformation

All scores were aligned such that higher scores indicated more difficulties. Thus, we reverse-coded (rev) VIQ, PIQ, LGC and PRC measures in ALSPAC and CC, FC and MR measures in ABCD. All measures were transformed to normality: First, scores were residualised for covariates. Rank-transformed residuals were again residualised for the same covariates to prevent the re-introduction of covariate effects (47). Covariates included age, age^2^, sex, sex×age, sex×age^2^, the first 10 ancestry-informative principal components from the genotyping analysis (EIGENSOFT, v6.1.4)(48) to correct for population stratification and, for ABCD only, genotyping batch effects.

### Multivariate phenotypic analyses

We performed phenotypic SEM (Supplementary methods) using exploratory factor analysis (EFA) and confirmatory factor analysis (CFA), based on split samples, each balanced for missingness, studying Pearson covariance matrices (r:base, pairwise-complete observations).

### Genome-wide genotyping information

<u>ALSPAC:</u> Genome-wide genotyping was performed using the Illumina HumanHap550 array (34,35). Standard quality control (QC) was applied as previously described (49) (Supplementary methods). In brief, 8,226 unrelated children (51% males) of European genetic ancestry (≤5526 with selected phenotype information) and 465,740 SNPs passed QC.

<u>ABCD:</u> Genome-wide information was obtained using the Affymetrix NIDA SmokeScreen Array (50) (Release 3.0) and QC applied (Supplementary methods). In total, 4,412 children (53% males) of European genetic ancestry (≤4,411 with selected phenotype information) and 398,864 SNPs passed QC.

### Genetic modelling

GRM construction: Genetic relationship matrices (GRMs) were calculated in ALSPAC and ABCD using PLINK, v1.9(51) based on directly genotyped genome-wide (genomic) markers with a relationship cut-off of 0.05, capturing the genetic relatedness among unrelated individuals.

Univariate and bivariate analyses of genetic influences: We assessed SNP-h² and r_g_ with univariate and bivariate genome-wide restricted maximum-likelihood (GREML), respectively, using Genome-wide Complex Trait Analysis (GCTA, v1.26.0) software (52,53), studying GRMs and phenotype information.

Multivariate modelling of genetic and residual structures: We model multivariate genomic and residual covariance structures by extending our previous genetic-relationship matrix structural equation modelling approach (GRM-SEM; (19,54,55)), allowing for r_GE_ as described below (R:grmsem, v1.1.6; https://gitlab.gwdg.de/beate.stpourcain/grmsem).

Here, we fitted multivariate models to jointly estimate the proportion of phenotypic variance attributable to genomic (SNP-h²) and residual influences (e²). Genetic (r_g_) and residual (r_e_) correlations between traits (or factors) measure the extent to which two phenotypes (or latent variables) share genetic or residual influences (range: −1 to 1). Using an extended modelling strategy, we also estimate correlations across genomic and residual factors (r_GE_, see below). Here, residual factors may reflect environmental contributions, but also systematic error or rare/non-additive genetic influences. As latter are either uncorrelated (56) or genetically distinct (57) from genomic factors, the presence of r_GE_ strengthens the evidence for environmental contributions captured by residual factors.

GRM-SEM dissects the phenotypic variance covariance into latent genomic and residual structures in samples of unrelated individuals, using GRMs. Analogous to GREML (53), the expected variance of a multivariate normal phenotype Y (Σ_V_) for 1..k traits is defined as the sum of the expected additive genetic (Σ_A_) and residual (Σ_E_) variance components where Σ_V_, Σ_A_, and Σ_E_ are symmetric k×k matrices.

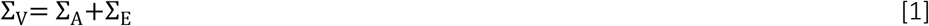

Using a maximum likelihood approach, genomic and residual variance are estimated with a structural model:

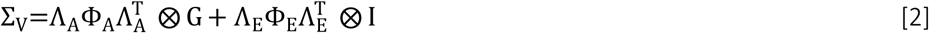

Λ_A_ and Λ_E_ capture genetic and residual factor loadings, and Φ_A_ and Φ_E_ genetic and residual factor (co)variance, with each factor variance constrained to one (i.e. a diagonal of 1). The dimensionality of Λ_A_ and Λ_E_, and Φ_A_ and Φ_E_ depends on the model, as described below and in the Supplement. Genetic-relatedness between pairs of n independent individuals is captured by G, a GRM matrix of n×n dimensions, while I is an n×n identity matrix. ⊗ indicates the Kronecker product.

*The Cholesky decomposition model* (Supplementary Figure 1a) fully parameterises both latent genetic and residual structures by fitting as many factors as observed phenotypes defining Λ_A_ and Λ_E_ as k×k lower triangular matrices, and Φ_A_ and Φ_E_ as k×k identity matrices (saturated model; Supplementary Figure 1a). Here, the Cholesky model is used as a descriptive baseline (saturated) model.

*The Independent Pathway (IP) model* jointly models latent genetic and residual structure (Figure 2a). For n_A_ genetic factors, genetic factor loadings are given by an augmented k×(n_A_+k) Λ_A_matrix, where the k×n_A_ part details genetic factor loadings and the k×k part details a diagonal matrix of specific genetic factor loadings (AS), with

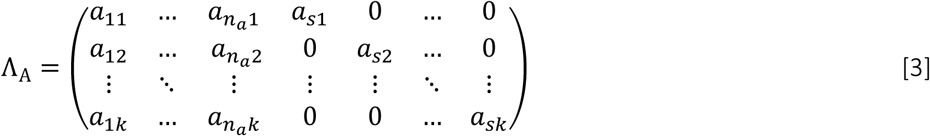

**Figure 2:**
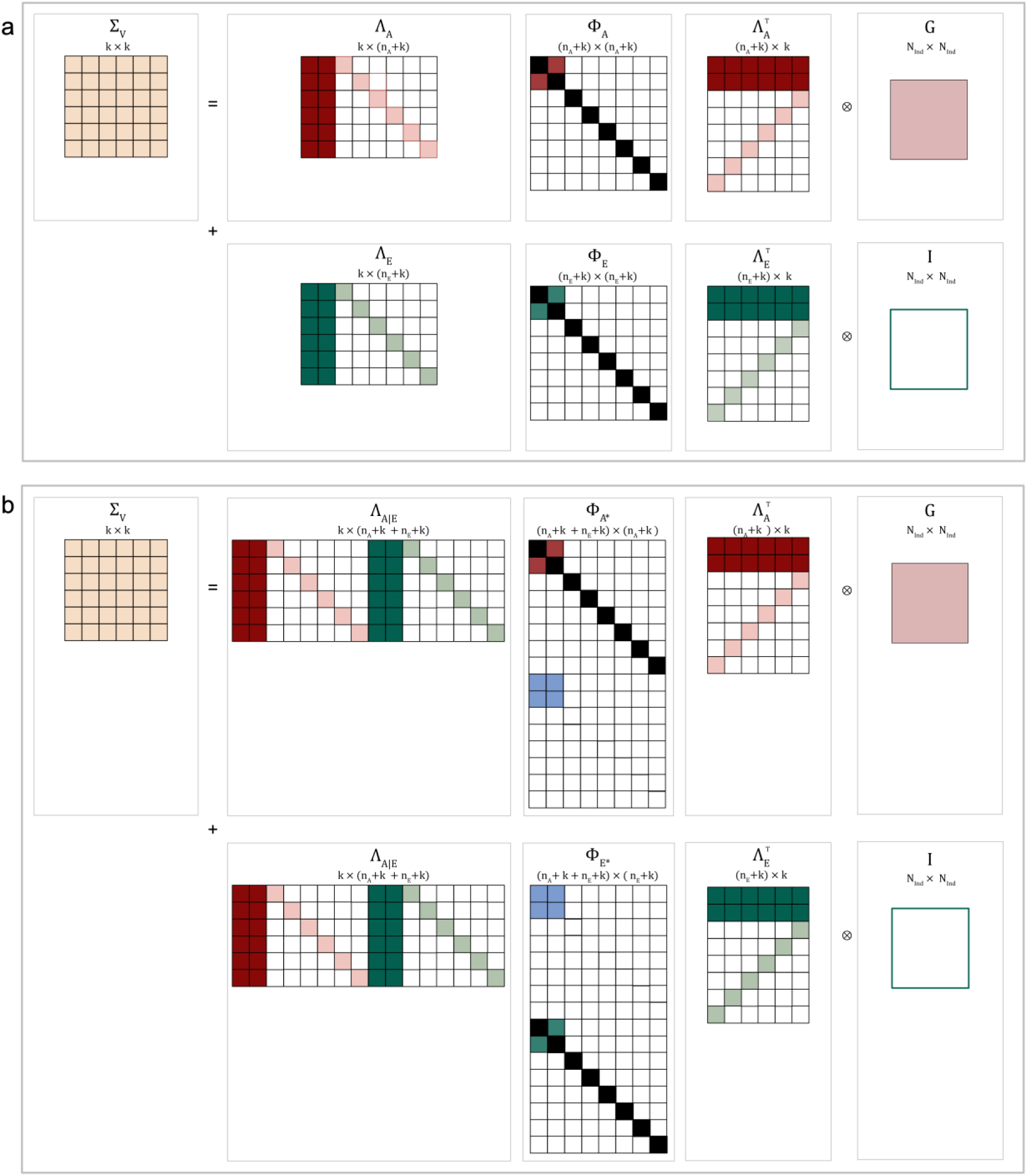
Schematic representation of GRM-SEM Independent Pathway models with and without r_GE_. Schematic representation of GRM-SEM model structures as illustrated for a six-variate trait shown for (a) an Independent Pathway model (with a 2-factor genetic and a 2-factor residual structure) without r_GE_ and (b) an Independent Pathway model (with a 2-factor genetic and a 2-factor residual structure) with r_GE_. For (a), Λ_A_ and Λ_E_ capture genetic and residual factor loadings, and Φ_A_ and Φ_E_ genetic and residual factor (co)variance, with each factor variance constrained to one (i.e. a diagonal of 1).For (b), Φ*_A_ and Φ*_E_ are extended Φ_A_ and Φ_E_ matrices including gene-environment covariance Φ_*cov*(*AE*)_. White squares indicate values of zero and black squares values of one. G – Genetic Relationship Matrix; GRM-SEM - Genetic-Relationship-Matrix Structural Equation Modelling; I – Identity matrix; k - Degrees of freedom; n_A_ - Number of genetic factor; n_E_ - Number of residual factors

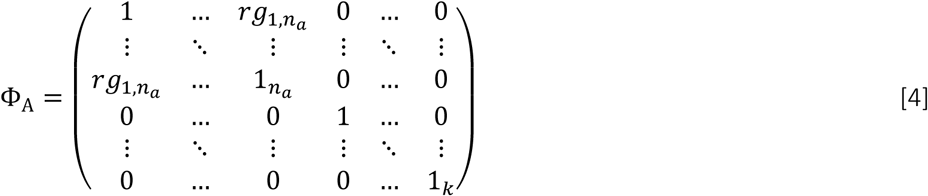

Consequently, Λ_A_is a k×(n_A_+k) matrix and Φ_A_ is a (n_A_+k)×(n_A_+k) matrix. Genetic factor correlations are estimated as off-diagonal elements in Φ_A_, specifically Φ_A_[1..n_A_,1..n_A_]. The residual part of the model is specified similarly, with Λ_E_ as a k×(n_E_+k) matrix of residual factor loadings and Φ_E_as a (n_E_+k)×(n_E_+k) matrix, where n_E_ is the number of residual factors (E) with k specific factors (ES), with

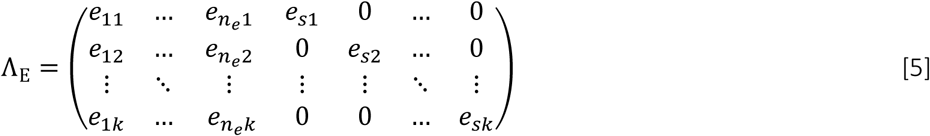

and

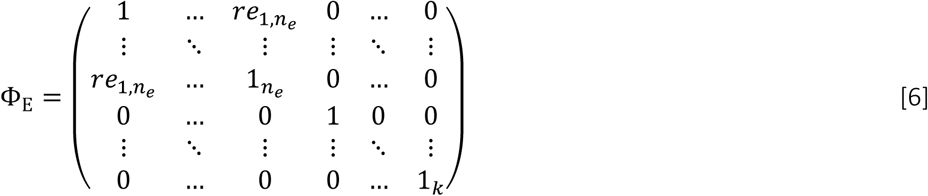

Residual factor correlations (r_e_) are estimated as off-diagonal elements in Φ_E_ capturing residual factors, i.e. Φ_E_[1..n_E_,1..n_E_].

Relaxing the assumption of independence of genetic and residual contributions, estimates of covariance across factors can be obtained by extending equation (2) as implemented for an IP model allowing for r_GE_ (Figure 2b). Adopting an augmented notation, the expected phenotypic variance can be described as

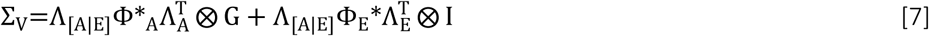

where Λ_[A|E]_ is an augmented matrix consisting of genetic (Λ_A_) and residual (Λ_E_) factor loadings, with k×(n_A_+k+n_E_+k) dimensions

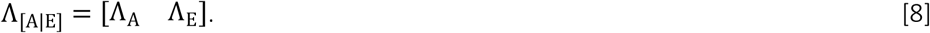

We assume that Φ is a hypothetical symmetric (n_A_+k+n_E_+k)×(n_A_+k+n_E_+k) correlation block matrix across all genetic and residual factors, with

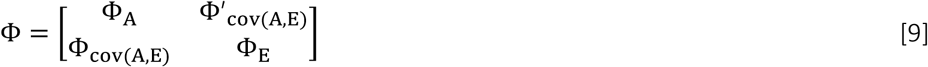

where Φ_cov(A,E)_ defines the (n_E_+k)×(n_A_+k) covariance matrix between genetic and residual factors loadings. Gene-environment correlations (r_GE_) are estimated between genetic and residual factors only, i.e. within Φ_cov(A,E)_[1.. n_E_,1.. n_A_], while correlations between specific genetic and specific residual factor loadings are absent, by definition, with

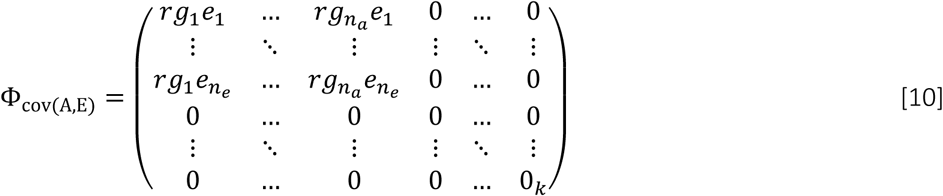

The symmetric correlation matrices Φ_A_ and Φ_E_ for genetic and residual factors have been described above. The diagonal of each matrix is fixed to one and serves here both as unit factors variance and as an indicator for the Kronecker product with G and I, respectively.

Thus, Φ can be understood as an augmented block matrix

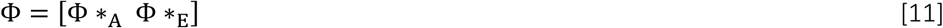

where

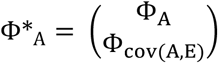

and

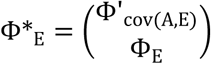

with Φ*_A_ being a 2(n_A_+k)×(n_A_+k) and Φ*_E_ a 2(n_E_+k)×(n_E_+k) matrix. Evidence for r_GE_ is obtained when the IP model allowing for r_GE_ fits the data better than the IP model assuming A and E independence, and the r_GE_ estimates are different from zero.

*The hybrid Independent Pathway/Cholesky (IPC) model* (Supplementary Figure 1b) dissects the phenotypic covariance into a latent variable structure, where the genetic part is captured with an Independent Pathway model, and the residual part with a Cholesky decomposition (see above).

*The hybrid Cholesky/Independent Pathway (CIP) model*, in turn, fully parameterises the genetic part using a Cholesky decomposition and the residual structure with an Independent Pathway model (Supplementary Figure 1c).

*Bifactor models* were fitted to test the independence of identified genetic and residual factors, respectively (Supplementary methods). A bifactor model defines one general factor loading on all traits, while any other additional factor defines the grouping of traits.

*Model fitting:* To accurately model multivariate genomic and residual structures, we adopted a data-driven analysis approach to define IPC/CIP and IP models informed by the Cholesky model and a combination of principal component analysis (PCA) and exploratory factor analysis (EFA) approaches (Figure 1), extending previous work focusing exclusively on genomic structures (30). We also derive the proportion of genetic or residual trait variance that can be explained by a latent genetic (factorial co-heritability, fc_SNP-h²_) or residual factor (factorial co-environmentality, fc_e²_), respectively.

### Model fit indices

The model fit of phenotypic and GRM-SEM models was assessed with AIC, BIC and SRMR (≤0.08), and, for phenotypic models only, incremental CFI, TLI (≥0.95) and RMSEA (≤0.06) indices (58).

## Results

### Study design

To disentangle phenotypic dimensions underlying cognition, language, and social skills in mid-childhood and early adolescence, we adopted a two-stage study design (Figure 1). We carried out discovery analyses in ALSPAC (stage 1) and followed-up findings in ABCD (stage 2) (Figure 1a), studying unrelated 8-13-year-old European-descent children from the general population (Supplementary Table 1). We identified genetic and residual structures across the selected measures by implementing a data-driven hybrid covariance modelling approach, including a combination of PCA, EFA and GRM-SEM techniques (Figure 1b). For the best-fitting model, we estimated r_GE_ across the identified genetic and residual factor structures (Figure 1c).

### Stage 1: Discovery analysis in ALSPAC

#### Covariance modelling without r_GE_

For multivariate modelling, we selected six of the 18 cognition, language, and social/social-communicated-related measures (age 8-13 years; Supplementary Table 1) with evidence for genomic contributions (*P*-value_GREML SNP-h²_<0.05; Supplementary Table 1, Supplementary Figure 2, Table 1): cognitive problems (PIQ_rev_9Y_: SNP-h²(SE)=0.24(0.066), VIQ_rev_9Y_: SNP-h²(SE)=0.53(0.065), language problems (LGC_rev_9Y_: SNP-h²(SE)=0.31(0.066)), social communication difficulties (SCD_8Y_: SNP-h²(SE)=0.17(0.065); PRC_rev_10Y_: SNP-h²(SE)=0.15(0.065)) and peer problems (PP_10Y_: SNP-h²(SE)=0.17(0.063).

**Table 1:**
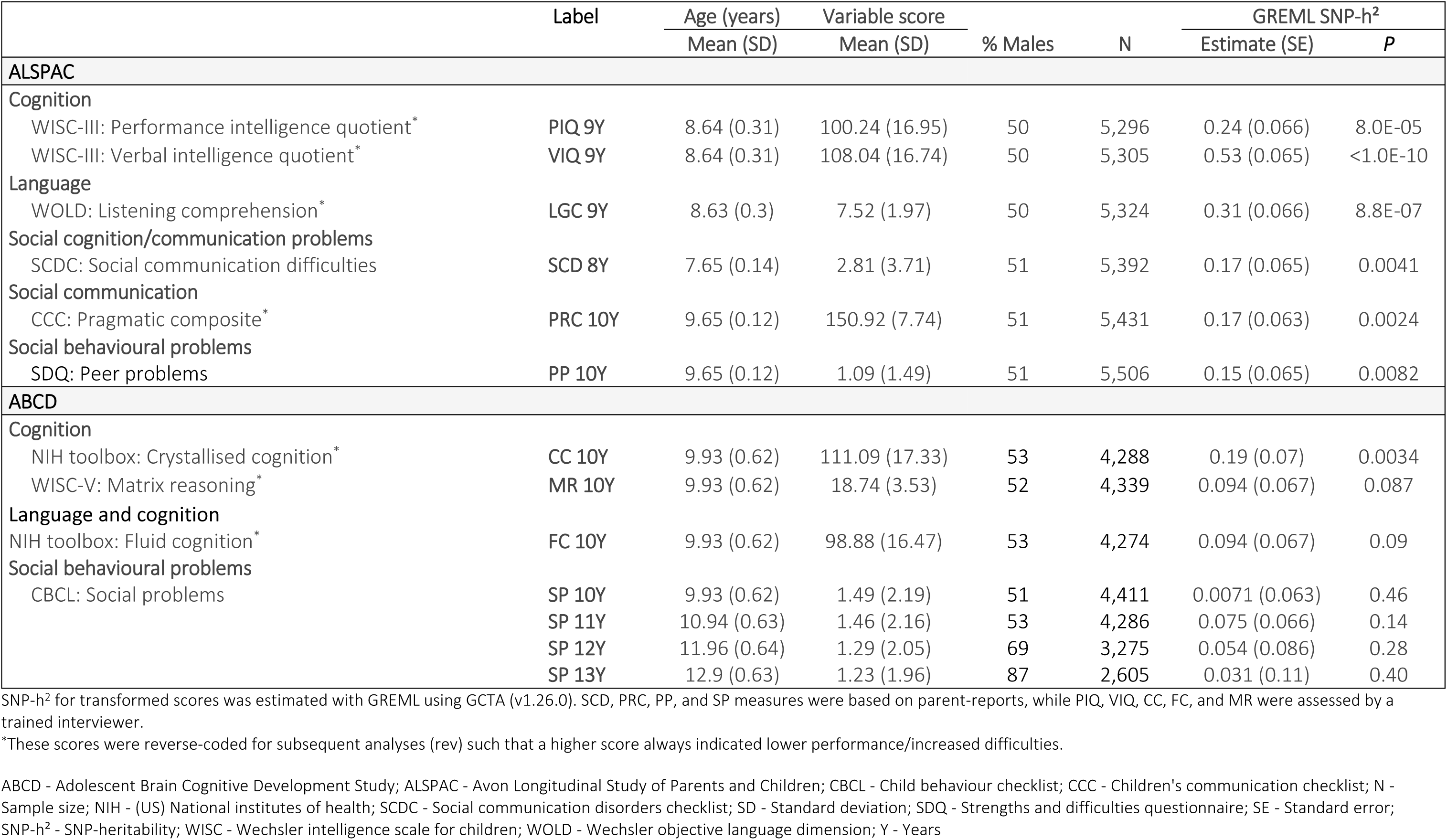
Descriptive information and SNP-h² estimates of cognitive and social/-communication traits in ALSPAC and ABCD.

|  | Label | Age (years) | Variable score | % Males | N | GREML SNP-h<sup>2</sup> |  |
| --- | --- | --- | --- | --- | --- | --- | --- |
|  |  | Mean (SD) | Mean (SD) |  |  | Estimate (SE) | P |
| ALSPAC |  |  |  |  |  |  |  |
| Cognition |  |  |  |  |  |  |  |
| WISC-III: Performance intelligence quotient* | PIQ 9Y | 8.64 (0.31) | 100.24 (16.95) | 50 | 5,296 | 0.24 (0.066) | 8.0E-05 |
| WISC-III: Verbal intelligence quotient* | VIQ 9Y | 8.64 (0.31) | 108.04 (16.74) | 50 | 5,305 | 0.53 (0.065) | <1.0E-10 |
| Language |  |  |  |  |  |  |  |
| WOLD: Listening comprehension* | LGC 9Y | 8.63 (0.3) | 7.52 (1.97) | 50 | 5,324 | 0.31 (0.066) | 8.8E-07 |
| Social cognition/communication problems |  |  |  |  |  |  |  |
| SCDC: Social communication difficulties | SCD 8Y | 7.65 (0.14) | 2.81 (3.71) | 51 | 5,392 | 0.17 (0.065) | 0.0041 |
| Social communication |  |  |  |  |  |  |  |
| CCC: Pragmatic composite* | PRC 10Y | 9.65 (0.12) | 150.92 (7.74) | 51 | 5,431 | 0.17 (0.063) | 0.0024 |
| Social behavioural problems |  |  |  |  |  |  |  |
| SDQ: Peer problems | PP 10Y | 9.65 (0.12) | 1.09 (1.49) | 51 | 5,506 | 0.15 (0.065) | 0.0082 |
| ABCD |  |  |  |  |  |  |  |
| Cognition |  |  |  |  |  |  |  |
| NIH toolbox: Crystallised cognition* | CC 10Y | 9.93 (0.62) | 111.09 (17.33) | 53 | 4,288 | 0.19 (0.07) | 0.0034 |
| WISC-V: Matrix reasoning* | MR 10Y | 9.93 (0.62) | 18.74 (3.53) | 52 | 4,339 | 0.094 (0.067) | 0.087 |
| Language and cognition |  |  |  |  |  |  |  |
| NIH toolbox: Fluid cognition* | FC 10Y | 9.93 (0.62) | 98.88 (16.47) | 53 | 4,274 | 0.094 (0.067) | 0.09 |
| Social behavioural problems |  |  |  |  |  |  |  |
| CBCL: Social problems | SP 10Y | 9.93 (0.62) | 1.49 (2.19) | 51 | 4,411 | 0.0071 (0.063) | 0.46 |
|  | SP 11Y | 10.94 (0.63) | 1.46 (2.16) | 53 | 4,286 | 0.075 (0.066) | 0.14 |
|  | SP 12Y | 11.96 (0.64) | 1.29 (2.05) | 69 | 3,275 | 0.054 (0.086) | 0.28 |
|  | SP 13Y | 12.9 (0.63) | 1.23 (1.96) | 87 | 2,605 | 0.031 (0.11) | 0.40 |

SNP-h<sup>2</sup> for transformed scores was estimated with GREML using GCTA (v1.26.0). SCD, PRC, PP, and SP measures were based on parent-reports, while PIQ, VIQ, CC, FC, and MR were assessed by a trained interviewer.
\*These scores were reverse-coded for subsequent analyses (rev) such that a higher score always indicated lower performance/increased difficulties.
ABCD - Adolescent Brain Cognitive Development Study; ALSPAC - Avon Longitudinal Study of Parents and Children; CBCL - Child behaviour checklist; CCC - Children's communication checklist; N - Sample size; NIH - (US) National institutes of health; SCDC - Social communication disorders checklist; SD - Standard deviation; SDQ - Strengths and difficulties questionnaire; SE - Standard error; SNP-h<sup>2</sup> - SNP-heritability; WISC - Wechsler intelligence scale for children; WOLD - Wechsler objective language dimension; Y - Years

We dissected the multivariate phenotypic covariance across the six measures into genomic and residual structures using GRM-SEMadopting a data-driven step-wise modelling approach, assuming independence of A and E influences (Figure 1b): First, we predicted the latent genetic structure with consecutive PCA (Figure 3a) and EFA (Supplementary Table 3) based on Cholesky-derived genetic correlation matrices, respectively, and identified a 2-factor IPC model (Supplementary Figure 3). Next, we predicted the latent residual structure from PCA (Figure 3a) and EFA (Supplementary Table 4) of the IPC-derived residual correlation and covariance matrices, respectively, to inform the model-building of an IP model.

**Figure 3:**
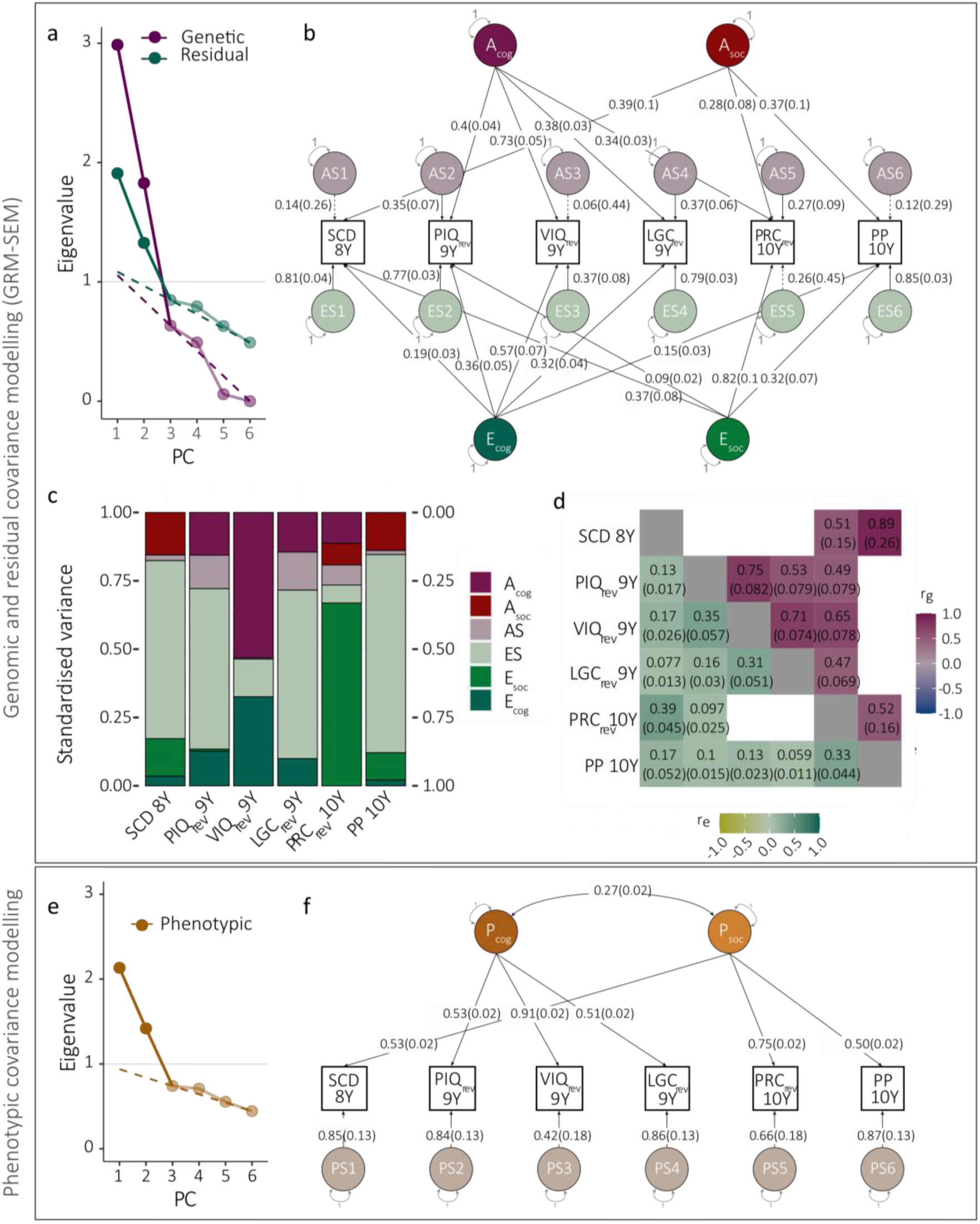
GRM-SEM and phenotype models in ALSPAC (without r_GE_). **(a)** Scree plot, **(b)** path diagram, **(c)** standardised genetic (h^2^) and non-genomic/residual (e^2^) variance, and **(d)** genetic (r_g_) and residual (r_e_) correlation patterns of the best fitting GRM-SEM (IP) model, without r_GE_ (N=6,543). **(e)** Scree plot and **(f)** path diagram of the best fitting phenotype model, based on a split-half approach (N=3,273). **(a,e)** The number of PCs was predicted using the Kaiser’s criterion (Eigenvalue>1; grey line), and Cattell’s scree test (dashed line). **(b,f)** Path diagrams with standardised factor loadings. The best-fitting GRM-SEM (IP) model (b) consists of genetic (A), specific genetic (AS), residual (E), and specific residual (ES) factors. The best-fitting phenotypic model (f) consists of phenotypic (P) and specific (PS) factors (specific factor loadings are shown as the square root of the specific factor variance). Observed measures are represented by squares and latent variables by circles. Single-headed arrows (paths) define relationships between variables, and double-headed arrows correlations. The variance of latent variables is constrained to unit variance. Note that SEs for GRM-SEM SNP-h^2^ contributions (c) have been omitted for clarity. ALSPAC - Avon Longitudinal Study of Parents and Children; IP - Independent pathway; LGC - Listening comprehension; PC - Principal component; PCA - Principal component analysis; PIQ - Performance intelligence quotient; PP - Peer problems; PRC - Pragmatic composite; rev – reverse-coded; SCD – social communication difficulties; VIQ - Verbal intelligence quotient; Y - Years

The model fit for the identified 2-factor GRM-SEM IP-model (Figure 3b-d) was comparable to the Cholesky model based on LRT (*P*_LRT_=0.19), but showed superior parsimony, and predicted the phenotypic variance/covariance well (SRMR=0.0071; Table 2, Supplementary Table 5). The factorial independence of both genetic and residual factors was confirmed with bifactor models (Table 2, Supplementary Table 5). The final GRM-SEM model included two independent genetic and two independent residual factors (Figure 3b-d; Supplementary Table 6), matching in structure a data-driven phenotypic model with modest factor correlation (r(SE)=0.27(0.016)) (Figure 3e,f, Table 2, Supplementary Table 7).

**Table 2:** Model fit comparison in ALSPAC and ABCD.

| ALSPAC ($N_{\text{traits}}=6$) | | | | | | | | | |
| --- | --- | --- | --- | --- | --- | --- | --- | --- | --- |
| GRM-SEM modelling ($N_{\text{ind}}=6,543$; $N_{\text{obs}}=32,254$) | | | | | | | | | |
| | Fig | $N_p$ | LL | AIC | BIC | SRMR | $\Delta df$ | LRT-$\Delta X^2$ | LRT-P |
| Cholesky |  | 42 | -13,397.26 | 26,878.51 | 27,163.53 | 0.0063 | - | - | - |
| IPC $n_A=2$ | S3b | 34 | -13,402.94 | 26,873.87 | 27,104.6 | 0.0061 | 8 | 11.36 | 0.18 |
| **IP $n_A=2$, $n_E=2$** | **3b** | **28** | **-13,406.46** | **26,868.92** | **27,058.94** | **0.0071** | **14** | **18.41** | **0.19** |
| IP $n_A=2$, $n_E=2$, $r_{GE}$ | | 30 | -13404.18 | 26,868.35 | 27,071.94 | 0.0071 | 12 | 13.84 | 0.31 |
| **IP $n_A=2$, $n_E=2$, $r_{GE \text{ con}}$** | **4a** | **28** | **-13404.18** | **26,866.35** | **27,054.37** | **0.0071** | **14** | **13.84** | **0.46** |
| Phenotypic modelling (EFA: $N_{\text{ind}}=3,273$; CFA: $N_{\text{ind}}=3,270$) | | | | | | | | | |
| | | $N_p$ | LL | AIC | BIC | SRMR | CFI | TLI | RMSEA |
| **$n_P=2$, correlated** | **3f** | **13** | **-31,647.23** | **63,320.46** | **63,402.16** | **0.023** | **0.99** | **0.98** | **0.037** |
| $n_P=2$ | | 15 | -31,648.57 | 63,325.14 | 63,413.12 | 0.034 | 0.99 | 0.97 | 0.041 |
| ABCD Study® ($N_{\text{traits}}=7$) | | | | | | | | | |
| GRM-SEM modelling ($N_{\text{ind}}=4,412$; $N_{\text{obs}}=27,478$) | | | | | | | | | |
| | | $N_p$ | LL | AIC | BIC | SRMR | $\Delta df$ | LRT-$\Delta X^2$ | LRT-P |
| Cholesky |  | 56 | -9,811.09 | 19,734.17 | 20,092.13 | 0.019 | - | - | - |
| **CIP $n_E=2$** | **5b** | **43** | **-9,818.95** | **19,723.90** | **19,998.76** | **0.020** | **13** | **15.73** | **0.26** |
| Phenotypic modelling (EFA: $N_{\text{ind}}=2,206$; CFA: $N_{\text{ind}}=2,206$) | | | | | | | | | |
| | | $N_p$ | LL | AIC | BIC | SRMR | CFI | TLI | RMSEA |
| **$n_P=2$, correlated** | **5f** | **15** | **-19575.63** | **39181.26** | **39266.74** | **0.027** | **0.98** | **0.97** | **0.050** |
| $n_P=2$ | | 16 | -19606.56 | 39181.13 | 39272.32 | 0.034 | 0.98 | 0.97 | 0.051 |

For phenotypic modelling, EFA and CFA were performed on split samples balanced for missingness. Phenotypic model fit was assessed with model fit parameters such as AIC, BIC, SRMR, CFI, TLI, and RMSEA.
ABCD - Adolescent Brain Cognitive Development Study; AIC - Akaike information criterion; ALSPAC - Avon Longitudinal Study of Parents and Children; BIC - Bayesian information criterion; con – Model with constrained parameters (constraining two near-zero specific genetic factor loadings); CFI - Comparative fit index; CIP - Hybrid Cholesky/independent pathway; df - Degrees of freedom; Fig – Corresponding figure or supplementary (S) figure; $n_A$ - Number of genomic factors; $n_E$ - Number of residual factors; IP - Independent pathway; IPC - Hybrid independent pathway/Cholesky; $N_{\text{ind}}$ - Number of individuals; $N_{\text{obs}}$ - Number of observations; $N_p$ - Number of parameters; $N_{\text{trait}}$ - Number of traits; P - Phenotypic factor; RMSEA - Root mean square error of approximation; SRMR - Standardised root mean square residual; TLI - Tucker-Lewis index

The first genetic factor (A_cog_) of the identified GRM-SEM IP model predominantly captured genetic variance (SNP-h²) in cognitive and language measures. The largest factor loading was found for VIQ scores (VIQ_rev_9Y_: λ_Acog_(SE)=0.73(0.052)), with the factor explaining nearly the entire SNP-h² (factorial co-heritability, fc_SNP-h²_(SE)=0.99(0.11), Supplementary Table 5). A_cog_ also explained genetic variance in PIQ (PIQ_rev_9Y_: λ_Acog_(SE)=0.40(0.044)), language comprehension (LGC_rev_9Y_: λ_Acog_(SE)=0.38(0.033)), and some social-communication difficulties (PRC_rev_9Y_: λ_Acog_(SE)=0.34(0.029)). The second genetic factor (A_soc_) accounted exclusively for variation in social and social-communication difficulties with the largest factor loading for SCDC scores (SCD_8Y_: λ_Asoc_(SE)=0.39(0.1)) and peer problems (PP_10Y_: λ_Asoc_(SE)=0.37(0.099)), capturing most of the SNP-h² (fc_SNP-h²_(SE)=0.88(0.41) and fc_SNP-h²_(SE)=0.90(0.45), respectively; Supplementary Table 6). A modest genetic factor loading was observed for pragmatic composite scores (PRC_rev_9Y_: λ_Asoc_(SE)=0.28(0.08)). The predicted GRM-SEM covariance structure was consistent with GREML-based SNP-h² and r_g_ estimates, based on overlapping 95% confidence intervals assuming multivariate normality (Supplementary Figure 4).

The residual structure of the identified IP model broadly mirrored the genomic structure. The first residual factor (E_cog_) captured residual variance across cognitive and language measures with the largest loading for VIQ (VIQ_rev_9Y_: λ_Ecog_(SE)=0.57(0.066)) explaining the majority of the standardised residual variance e^2^. In other words, the proportion of e² in VIQ explained by λ_Ecog_ was large (factorial co-environmentality, fc_e²_(SE)=0.70(0.12); Supplementary Table 6). The second residual factor (E_soc_) explained residual variance across all social scores, with the largest factor loading for the pragmatic composite (PRC_rev_9Y_: λ_Esoc_(SE)=0.82(0.14), fc_e²_(SE)=0.91(0.31), Supplementary Table 6). However, cross-loadings were negligible (λ≤0.3) in contrast to the genomic factor structure.

#### Covariance modelling with r_GE_

Given the similarity in phenotypic, genomic and residual structures, we subsequently studied the correlation across genomic and residual factors (r_GE_), both for the cognitive (r_GE(cog)_: A_cog_, E_cog_) and the social dimension (r_GE(soc)_: A_soc_, E_soc_)(Figure 4). Specifically, we extended the best-fitting GRM-SEM IP, assuming A and E independence, model with two r_GE_ parameters and re-fitted the model (Figure 4a, Supplementary Table 5, Supplementary Table 6). Once parameters explaining little variance (λ_AS_<0.05) were constrained to zero, the IP_r_GE_ model fitted the data best (Table ^2^).

**Figure 4:**
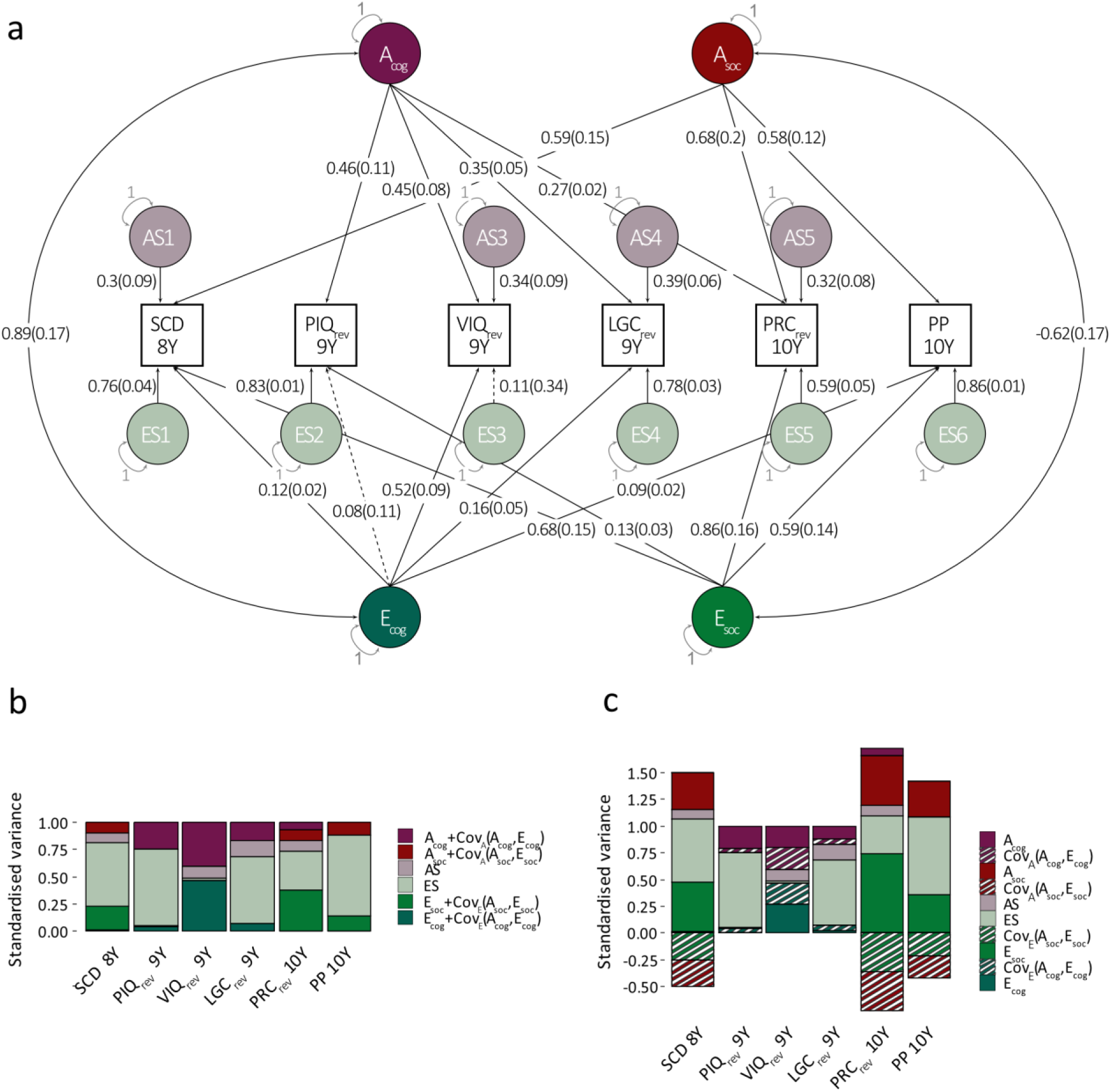
GRM-SEM model in ALSPAC (with r_GE_). **(a)** Path diagram with standardised factor loadings and corresponding standard errors based for the best-fitting GRM-SEM IP model allowing for r_GE_ (N=6,543). Here, the best-fitting IP model assuming the independence of A and E (Figure 3b) was extended with r_GE_ parameters, and re-fitted. **(b,c)** Different representations of the same standardised genetic and residual variance contributions from model (a): **(b)** Combined standardised genetic and residual variance contributions, including A, Cov(A,E) (G related), as well as E and Cov(A,E) (E-related) contributions. **(c)** Individual standardised genetic and residual variance contributions shown for A, E and 2Cov(A,E) (i.e. G and E-related), respectively, allowing for different directions of effect. The GRM-SEM IP model consists of genetic (A), specific genetic (AS), residual (E), and specific residual (ES) factors, as well as the covariance between genetic and residual factors, i.e. Cov(A,E) either with respect to genetic (striped red) or residual (striped green) contributions. Observed measures are represented by squares and latent variables by circles. Single-headed arrows (paths) define relationships between variables, and double-headed arrows correlations. The variance of latent variables is constrained to unit variance. Estimates in panels (b) and (c) were derived from the model in (a). Note that SEs for GRM-SEM variance contributions (b,c) have been omitted for clarity. ALSPAC - Avon Longitudinal Study of Parents and Children; E – Non-genomic/residual factors; r_GE_ – Gene-environment correlation; IP - Independent pathway; LGC - Listening comprehension (Wechsler objective language dimension); PIQ - Performance intelligence quotient (Wechsler intelligence scale for children III); PP - Peer problems (Strengths-and-difficulties questionnaire); PRC - Pragmatic composite scores (Children’s communication checklist); rev – reverse-coded; SCD – social communication difficulties (Social communication disorders checklist); VIQ - Verbal intelligence quotient (Wechsler intelligence scale for children III); Y - Years

There was positive r_GE_ across the cognitive/language dimension (r_GE(cog)_=0.89(SE=0.18), *P*=8.8 ×10^−6^). Compared to the unadjusted model, genetic and residual factor loadings for cognitive measures decreased, most strongly for VIQ genetic loadings (IP model VIQ_rev_9Y_: λ_Acog_=0.73(SE=0.05) versus IP_r_GE_ model VIQ_rev_9Y_: λ_Acog_=0.45(SE=0.08)). This implies that positive covariance between A and E (Cov_A_(A_cog_, E_cog_), Figure 4c) adds to the true genetic variance for cognitive measures including PIQ_rev_, VIQ_rev_ and LGC_rev_ (A_cog_, Figure 4c), increasing measurable genetic variance contributions (A_cog_+Cov_A_(A_cog_, E_cog_), Figure 4b). In analogy, the positive covariance between A and E also inflates the measurable residual variance (E_cog_, Cov_E_(A_cog_, E_cog_), Figure 4c; E_cog_+Cov_E_(A_cog_, E_cog_), Figure 4b).

In contrast, r_GE_ for the social (problem) dimension was negative (r_GE(social)_=-0.62(SE=0.17)), *P*=5.0×10^−4^), consistent with compensatory mechanisms. Compared to the unadjusted IP model, IP_r_GE_ genetic and residual factor loadings for social phenotypes increased in value, although 95% CI bands overlapped throughout. Thus, negative covariance between A and E (Cov_A_(A_soc_, E_soc_), Figure 4c) subtracts from the true genetic variance for social measures such as SCD, PRC_rev_ and PP scores(A_soc_, Figure 4c), resulting in lower measurable genetic variance contributions (A_soc_+Cov_A_(A_soc_, E_soc_), Figure 4b). Residual variances showed a similar pattern (E_soc_, Cov_E_(A_cog_, E_cog_), Figure 4c; E_cog_+Cov_E_(A_cog_, E_cog_), Figure 4b).

Thus, genomic and residual structures for cognitive versus social traits are distinct and trait-specific r_GE_, consistent with different aetiological mechanisms underlying each phenotypic domain.

### Stage 2: Follow-up analysis in ABCD

#### Covariance modelling without r_GE_

We followed up our findings in ABCD, investigating cognitive, language and social measures at ages 10 to 13 years (Supplementary Table 2; note that social communication assessments were not available). While data were collected using different psychological instruments, the underlying concepts were broadly consistent across studies. However, across the selected measures there was little evidence for SNP-h² (Supplementary Figure 5, Supplementary Figure 6a). Consequently, we studied the latent residual structure using a hybrid Cholesky/Independent Pathway (CIP) model (Figure 5, Supplementary Tables 8-9).

**Figure 5:**
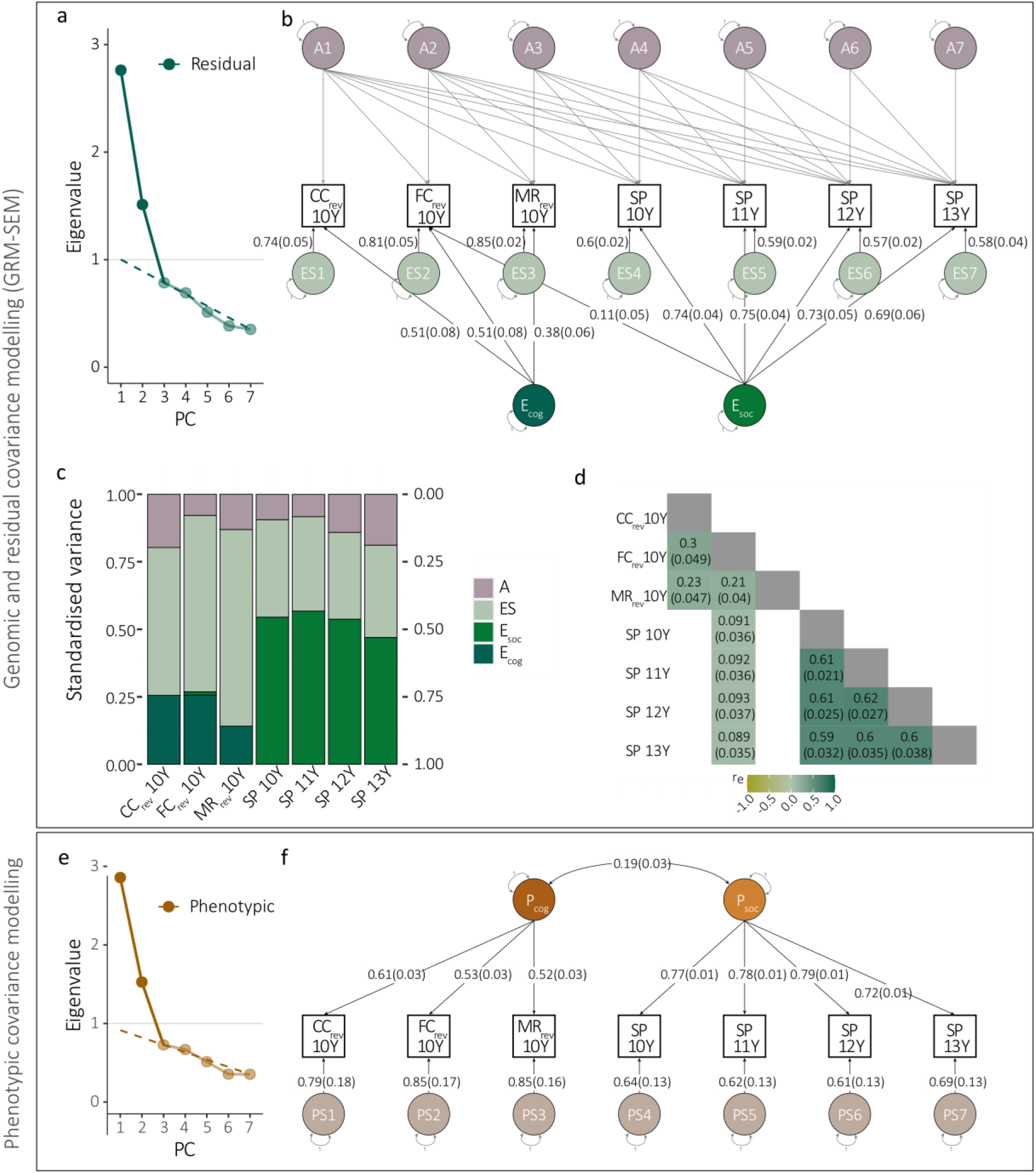
GRM-SEM and phenotype models in ABCD (without r_GE_). **(a)** Scree plot, **(b)** path diagram, **(c)** standardised genetic and residual variance, and **(d)** genetic and residual correlation patterns of the best fitting GRM-SEM model (CIP) (N=4,412). **(e)** Scree plot and **(f)** path diagram of the best fitting phenotype model, based on a split-half approach (N=2,206). **(a,e)** The number of PCs was predicted with PCAs using the Kaiser’s criterion (Eigenvalue>1; grey line), and Cattell’s scree test (dashed line). **(b,f)** Path diagrams with standardised factor loadings and corresponding standard errors for the best-fitting models. The best-fitting GRM-SEM (CIP) model (b) consists of genetic (A), residual (E), and specific residual (ES) factors. The best-fitting phenotypic model (f) consists of phenotypic (P) and specific (PS) factors (specific factor loadings are shown as the square root of the specific factor variance). Observed measures are represented by squares and latent variables by circles. Single-headed arrows (paths) define relationships between variables, and double-headed arrows correlations. The variance of latent variables is constrained to unit variance. Note that SEs for GRM-SEM SNP-h^2^ contributions (c) have been omitted for clarity. Due to little evidence for trait SNP-h², as estimated with GREML, we only modelled the latent residual structure in ABCD using a CIP model. ABCD - Adolescent Brain Cognitive Development Study; CC - Crystalized cognition (NIH Toolbox); CIP - Hybrid Cholesky/Independent pathway; FC - Fluid cognition; GREML – Genomic Restricted maximum likelihood; MR - Matrix reasoning; SNP-h² - SNP-heritability; SP - Social problems; Y - Years

Consistent with two predicted dimensions (Figure 5a), we identified a 2-factor CIP model (Figure 5b-d, Supplementary Table 9) with excellent model fit (Table 2, Supplementary Table 10). The CIP model included two uncorrelated residual factors, as confirmed using bifactor models (see Supplementary Table 10). The two-dimensional residual factor structure closely matched the two-dimensional structure of a data-driven phenotypic model (Figure 5e-f, Table 2, Supplementary Table 11) and the phenotypic structure previously observed in ALSPAC: E_cog_ accounted for residual variance underlying cognition and language with the largest factor loading for reverse-coded crystallised cognition (CC_rev_10Y_: λ_Ecog_(SE)=0.51(0.082), fc_e²_(SE)=0.32(0.094)), while E_soc_ explained residual variance across social traits with the largest loading for social problems (SP_11Y_: λ_Esoc_(SE)=0.75(0.04), fc_e²_(SE)=0.62(0.032)). The CIP-predicted covariance structure was consistent with GREML-based SNP-h² and r_g_ estimates (Supplementary Figure 6). Thus, the 2-factor phenotypic and residual structure in ABCD matched the 2-factor genetic, residual and phenotypic structure in ALSPAC.

## Discussion

Studying population-based cognitive, language, and social abilities in ALSPAC and ABCD using a hybrid covariance modelling approach, we observed structural differences between a cognitive/language versus a social domain, with striking similarities across genomic, phenotypic and/or residual influences within each domain. Trait-specific r_GE_ (ALSPAC only) manifested as a positive link across the cognitive/language domain and a negative link across the social domain.

The identified cognitive/language and social domains involved, for heritable traits, a combination of A and E factors (ALSAPC), and otherwise E only (ABCD). The majority of measures mapped to one phenotypic domain only (factor loading: λ≥0.3), suggesting distinct developmental mechanisms. These findings are consistent with neuro-imaging and behavioural research suggesting different neuro-cognitive mechanisms for social versus non-social learning (59,60). For example, fMRI studies have identified brain activation patterns linked to high-level language processing that remain unresponsive to social stimuli such as action observation and action imitation (61). In addition, the identification of an independent social factor is consistent with genetic research suggesting a non-cognitive domain of educational attainment that is uncorrelated with the cognitive domain (62). Cross-loadings across the cognitive and social factor were exclusively detected for pragmatic communication in ALSPAC at the genetic level, suggesting that during specific cognitive tasks, such as listening comprehension, cognitive/language and social brain-networks may synchronise (63), consistent with a recently proposed theoretical framework for human communication (64).

The identification of r_GE_ across pairs of structurally similar A and E factors (ALSPAC only) that differentiate cognitive/language from social domains demonstrates that modelling shared residual variation can provide meaningful insight into underlying aetiological mechanisms. Given r_GE_, the identified shared residual structures are inconsistent with systematic error or rare-variant contributions (Methods). The detection of r_GE_ in cohorts of unrelated individuals, excluding within-family effects, therefore suggests that shared residual factors are capturing higher-order environmental influences shared across families, such as regional influences, including neighbourhoods and school districts or genetic migration patterns across geographical regions (26,65).

The positive nature of r_GE_ for cognitive/language indicates an overestimation of both genetic and residual variance, here mostly strongly for verbal IQ, in unadjusted models. Our findings are consistent with reports of positive r_GE_ for traits such as educational attainment, household income and fluid intelligence in relation to schools, between-family variation and geographical region (26,66,67) involving mechanisms of “genomic nurture” (27). In contrast, negative r_GE_ for the social (problem) dimension suggests an underestimation of genetic and residual variance in genomic studies, as variance is cancelled out due to negative GE covariance. For example, children with social problems might have been given additional opportunities (e.g. by their parents or schools) to engage in positive social interactions, given that many schools aim to reduce social behavioural problems with preventive programs (68), consistent with evocative mechanisms (69).

This study has multiple strengths: First, we implemented a novel hybrid covariance modelling approach. Here, we jointly model genomic and residual structures and their r_GE_ in population-based cohorts, without relying on family-based structures. Second, our study comprised a broad set of well-established child/early-adolescent measures that capture variability in cognitive, language and social traits, although other questionnaires across different developmental periods may reveal different structures. Third, our findings are supported by two independent population-based cohorts from the UK and the US. We replicated both residual and phenotypic structures in ALSPAC and ABCD with matching genomic structures in ALSPAC, underlining the robustness of our findings. However, this study also has limitations. The lack of SNP-h^2^ for cognitive, language and social measures in ABCD, compared to ALSPAC, suggests differences in aetiological mechanisms and/or sample composition. For example, in ABCD the diversity in recruited population strata (70) might affect the power to detect genetic variance and r_GE_ by enhancing heterogeneity. In addition, analyses have been exclusively conducted across samples of European descent. Cognitive and social abilities may vary across different cultures (71), which, if linked to differences in genetic ancestries, can result in confounding, requiring future multi-ancestry studies.

In conclusion, by implementing a hybrid genomic and residual covariance modelling approach, we show that cognitive/language abilities versus social abilities are related to distinct phenotypic, residual and/or genomic contributions, including trait-specific r_GE_, suggesting, in childhood, differences in developmental mechanisms.

## Acknowledgements

We are extremely grateful to all the ALSPAC families who took part in this study, the midwives for their help in recruiting them, and the whole ALSPAC team, which includes interviewers, computer and laboratory technicians, clerical workers, research scientists, volunteers, managers, receptionists and nurses. The UK Medical Research Council and Wellcome (Grant ref: 217065/Z/19/Z) and the University of Bristol provide core support for ALSPAC. This publication is the work of the authors and they will serve as guarantors for the contents of this paper. A comprehensive list of grants funding is available on the ALSPAC website (http://www.bristol.ac.uk/alspac/external/documents/grant-acknowledgements.pdf). ALSPAC GWAS data was generated by Sample Logistics and Genotyping Facilities at Wellcome Sanger Institute and LabCorp (Laboratory Corporation of America) using support from 23andMe. BSTP and SEF are supported by the Max Planck Society. BSTP is also supported by R2D2-MH, which has been funded by Horizon Europe (grant agreement no. 101057385), by UK Research and Innovation (UKRI) under the UK government’s Horizon Europe funding guarantee [grant no. 10039383] and by the Swiss State Secretariat for Education, Research and Innovation (SERI) under contract number 22.00277

Data used in the preparation of this article were obtained from the Adolescent Brain Cognitive Development℠ (ABCD) Study (https://abcdstudy.org), held in the NIMH Data Archive (NDA). This is a multisite, longitudinal study designed to recruit more than 10,000 children age 9-10 and follow them over 10 years into early adulthood. The ABCD Study® is supported by the National Institutes of Health and additional federal partners under award numbers U01DA041048, U01DA050989, U01DA051016, U01DA041022, U01DA051018, U01DA051037, U01DA050987, U01DA041174, U01DA041106, U01DA041117, U01DA041028, U01DA041134, U01DA050988, U01DA051039, U01DA041156, U01DA041025, U01DA041120, U01DA051038, U01DA041148, U01DA041093, U01DA041089, U24DA041123, U24DA041147. A full list of supporters is available at https://abcdstudy.org/federal-partners.html. A listing of participating sites and a complete listing of the study investigators can be found at https://abcdstudy.org/consortium_members/. ABCD consortium investigators designed and implemented the study and/or provided data but did not necessarily participate in the analysis or writing of this report. This manuscript reflects the views of the authors and may not reflect the opinions or views of the NIH or ABCD consortium investigators.

ABCD data repository grows and changes over time The ABCD data used in this study came from the ABCD Data Release 3.0 (genotype data) and 4.0 (phenotype data) (https://doi.org/10.15154/1523041).

