## Supplementary Material for "Disentangling multivariate relationships between cognition, language and social traits: structures of G, E, and r_GE_"

#### Supplementary Methods

*Description of the ALSPAC cohort*

Pregnant women resident in Avon, UK with expected dates of delivery between 1st April 1991 and 31st December 1992 were invited to take part in the study (1,2). 20,248 pregnancies have been identified as being eligible and the initial number of pregnancies enrolled was 14,541. Of the initial pregnancies, there was a total of 14,676 foetuses, resulting in 14,062 live births and 13,988 children who were alive at 1 year of age. When the oldest children were approximately 7 years of age, an attempt was made to bolster the initial sample with eligible cases who had failed to join the study originally. As a result, when considering variables collected from the age of seven onwards (and potentially abstracted from obstetric notes) there are data available for more than the 14,541 pregnancies mentioned above: The number of new pregnancies not in the initial sample (known as Phase I enrolment) that are currently represented in the released data and reflecting enrolment status at the age of 24 is 906, resulting in an additional 913 children being enrolled (456, 262 and 195 recruited during Phases II, III and IV respectively). The phases of enrolment are described in more detail in the cohort profile paper and its update (see footnote 5 below). The total sample size for analyses using any data collected after the age of seven is therefore 15,447 pregnancies, resulting in 15,658 foetuses. Of these 14,901 children were alive at 1 year of age. Of the original 14,541 initial pregnancies, 338 were from a woman who had already enrolled with a previous pregnancy, meaning 14,203 unique mothers were initially enrolled in the study. As a result of the additional phases of recruitment, a further 630 women who did not enrol originally have provided data since their child was 7 years of age. This provides a total of 14,833 unique women (G0 mothers) enrolled in ALSPAC as of September 2021. G0 partners were invited to complete questionnaires by the mothers at the start of the study and they were not formally enrolled at that time. 12,113 G0 partners have been in contact with the study by providing data and/or formally enrolling when this started in 2010. 3,807 G0 partners are currently enrolled.

*Cognitive, language and social measures in ALSPAC*

*Cognition:*

A short form of the Wechsler Intelligence Scale for Children (WISC-III)(3), consisting of alternate items for all subtests except for the coding subtest, was administered to measure children's *Verbal and performance intelligence scores* at age 8 (WISC-III, VIQ, PIQ). The WISC-III includes ten subtests. Five subtests are verbal: information, similarities, arithmetic, vocabulary, comprehension, and can be used to construct a verbal intelligence score. The other five subtests are performance subtests: picture completion, coding, picture arrangement, block design and object assembly. Raw scores were calculated using items used in the alternate item form of the WISC-III. Total age-scaled verbal and performance scores were calculated using the look-up tables provided in the WISC-III manual, with a maximum VIQ score of 160 and a maximum PIQ score of 160. All scores were pro-rated. Test-retest correlations of the WISC-III verbal intelligence are high and range between 0.90 and 0.94, depending on the age at assessment and the duration of the test-retest interval (4). The VIQ is also highly correlated with the Kaufman Brief Intelligence Test (0.79) and with the Standford-Binet IV (0.69)(4). Test re-test correlations for the WISC-III performance intelligence are 0.89 (5). The correlation between PIQ assessed using WISC-III and the non-verbal score measured using the Otis-Lennon School Ability Test was 0.59 (6).

*Language:*

Children's listening comprehension (LGC) was measured using a subtest of the Wechsler Objective Language Dimensions(7) at age 8. The child was shown a picture while the tester read a paragraph about the image aloud. Next, the child was asked to answer fifteen questions about what they had heard. A listening comprehension score was calculated as the sum of correct answers. The LGC subtest has a test-retest reliability of 0.83 to 0.88 for children aged six to eleven years (8) and is correlated 0.44 (9) with the Peabody Picture Vocabulary Test-III (10).

*Social traits:*

Measures of social communication *difficulties* (SCD) at 8 and 11 years were obtained with the 12-item Social-Communication Disorder Checklist (SCDC; score range: 0 to 24)(11), using parent reports. The SCDC is a brief screening instrument of social reciprocity and verbal/nonverbal communication (e.g. "Not aware of other people's feelings"), with high reliability and good validity (11), which has been extensively investigated (11–13). Higher SCDC scores reflect more social-communication deficits.

Children's *social communication* abilities at 10 years were assessed with parent-reported pragmatic composite scores (PRC) using the Children's Communication Checklist (CCC) (14). The pragmatic summary score comprises the CCC subscales C to G: (C) inappropriate initiation, (D) coherence, (E) stereotyped conversation, (F) use of conversational context, and (G) conversational rapport. CCC subscales have a moderate/high consistency (0.62-0.83) and high reliability (0.74-0.87)(15).

*Social behaviour* was captured with the prosocial behaviour (PB) and peer problems (PP) subscales of the Strengths and Difficulties Questionnaire (SDQ)(16) and measured at the ages of 8, 10, 12, and 13 years based on parent reports, and at the ages of 9 and 11 years based on teacher reports. The reliability of the parent-reported SDQ PB and PP scales is sufficient (internal consistency as measured by Cronbach's $\alpha$ is 0.57 for the PP and 0.65 for the PB scale) (17). The validity of the SDQ has been assessed by how strongly the subscales are associated with psychiatric conditions (17), and high SDQ problem scores have been associated with a substantial increase in psychiatric risk. The peer problem subscale includes the five items: (I) "Rather solitary, tends to play alone"; (II) "Has at least one good friend"; (III) "Generally liked by other children"; (IV) "Picked on or bullied by other children"; and (V) "Gets on better with adults than with other children". Items II and III were reverse-coded, and eventually, all items were summed up to give a final peer problem score (score-range 0–10) with higher scores reflecting more peer-related problems. The prosocial scale includes the five items: " Considerate of other people's feelings "; "Shares readily with other children (treats, toys, pencils, etc.)"; "Helpful if someone is hurt, upset or feeling ill"; "Kind to younger children"; and "Often volunteers to help others (parents, teachers, other children)". All items were summed up to give a final prosocial score (score range 0–10), with higher scores reflecting more prosocial behaviour.

*<u>Genotype cleaning of the ALSPAC cohort</u>*

ALSPAC children were genotyped using the Illumina HumanHap550 quad chip, and genotypes were called using the Illumina GenomeStudio software. Standard genomic quality control (QC) (18) was performed at both the SNP and individual level using PLINK (v1.07)(19). Individuals with a sex mismatch, with single nucleotide polymorphism (SNP) missingness >3%, non-European ancestry and a genetic relationship >0.05 were excluded from the study. SNPs with a low call rate (<99%), low minor allele frequency (MAF<1%) or deviations from Hardy-Weinberg equilibrium ($P<5x10^{-7}$) were excluded, too. After QC, 8,226 unrelated children (51% males) of European genetic ancestry and 465,740 SNPs remained. Among those individuals, up to 5,516 children had both phenotypic and genome-wide information available and were analysed in this study.

*Description of the ABCD Study®*

The ABCD Study® is a US population-based longitudinal study of brain development and child health (20). Children were enrolled into the study at the age of 9-10 years. ABCD Study participants were followed for 10 years at 21 data acquisition sites across the US (N=11,877, Data Release 4.0 (21)). Participants were recruited through school systems, sampling across gender, race and ethnicity, socioeconomic status, and urbanicity (22). The Institutional Review Board (IRB) at the University of California, San Diego, approved all aspects of the ABCD Study (23). Parents or guardians provided written consent, while children provided written assent (24).

ABCD Study participants were recruited across diverse population strata (22). In particular, for each of the 21 study sites, the ABCD study employed a probability sampling strategy to identify schools within the catchment areas as the primary method for contacting and recruiting eligible children and their parents (25). Note that we did not apply ABCD sampling weights as the impact of population weights in genetic analyses is not yet fully understood, even within the ABCD (25), and may, besides unbiasing genetic association findings (26), also increase reporting error (27).

*Cognitive, language and social measures in ABCD*

*Cognition and Language:*

For this study, we analysed Cognition Fluid Composite and Crystallized Composite age-corrected standard scores from the ABCD Youth NIH Toolbox. The NIH Toolbox (http://www.nihtoolbox.org) is a testing battery developed to assess cognition measures within the NIH Blueprint for Neuroscience Research. It consists of seven tasks assessing episodic memory, executive function, attention, working memory, processing speed, and language abilities. Age correction was based on normative scores from a sample of 2,917 children and adolescents (28).

The derived Crystallized Intelligence Composite Score was based on the Toolbox Picture Vocabulary Task®(TPVT) and the Toolbox Oral Reading Recognition Task®(TORRT)(29,30). The TPVT measures language and verbal intellect. During the task, children have to match words listened to in audio files to pictures presented depicting the concept, idea or object referenced by the audio. The TORRT is a reading test where children have to pronounce single letters or words presented on an iPad and are evaluated by a trained tester.

The Fluid Intelligence Composite was based on Toolbox Pattern Comparison Processing Speed Test® (TPCPST)(31–33), the Toolbox List Sorting Working Memory Test® (TLSWMT)(34,35), the Toolbox Picture Sequence Memory Test® (TPSMT)(36,37), Toolbox Flanker Task® (TFT)(38,39), and the Toolbox Dimensional Change Card Sort Task® (TDCCS)(38–40). The TPCPST measures rapid visual processing speed during a task where children are shown two pictures and are asked to indicate whether the presented pictures are the same. The TLSWMT is a picture-based working memory task where participants have to sequence stimuli based on category membership and perceptual stimuli. The TPSMT assesses episodic memory based on a task where the participant has to retrieve the sequence of a series of fifteen pictures presented to the participant in a non-speeded setting. The TFT assesses executive function, attention and inhibition abilities based on a task where children have to indicate the direction of a stimulus while being presented congruent and incongruent secondary stimuli. Participants are evaluated based on speed and accuracy. The TDCCS measures executive function and cognitive flexibility within a sorting task where an object has to be matched to one of two other objects.

*Social behaviour*

Social behaviour was measured with the 11-item social problems (SP) subscale of the Child Behaviour Check List (41), which is part of the Achenbach System of Empirically Based Assessment (ASEBA), at 10, 11, 12, and 13 years, based on parent reports (42). The CBCL comprises 113 items assessing a broad range of behavioural and emotional problems. The SP scale assesses problems related to social functioning, such as difficulties with peer relationships, lack of social skills, and little adaptation to social contexts, such as "Not liked by other kids", "Prefers to be alone", or "Picked on or bullied". The parent-reported SP scale has a sufficient internal consistency (Cronbach's α ~0.65)(41). The summed raw scores are typically converted into standardised T-scores, based on age and gender-adjusted normative data. However, here we studied SP raw scores adjusted for age and sex, given that the available population-based sample size is considerably larger (>4,000 children) capturing a defined developmental stage (here mid-childhood to adolescence) compared to the original ASEBA normative data (1,753 children, aged 6 to 18 years).

*Genotype cleaning of the ABCD cohort*

ABCD children were genotyped with the Affymetrix NIDA SmokeScreen Array (43) (Release 3.0. Standard genomic quality control (18) was performed at both the SNP and individual level using PLINK (v1.9.0). Individuals with sex mismatches, with SNP missingness >3%, non-European ancestry, heterozygosity outliers (> 3SD) and a genetic relationship >0.05 were excluded from the study. SNPs with a low call rate (<99%), a MAF <1% or deviations from Hardy-Weinberg equilibrium ($P<5x10^{-7}$) were removed, too. In total, 4,475 children of European ancestry (53% males) and 398,864 SNPs passed QC. Of these individuals, 4411 had selected phenotype information and were included in this study.

*Principal component analysis (PCA)*

PCA was performed to identify the minimum number of dimensions required to describe the genetic, residual and phenotypic covariance structure. Eigenvalues were calculated based on genetic and residual correlation matrices, which were estimated using Cholesky decompositions in GRM-SEM. Eigenvalues were then evaluated according to Kaiser's rule, and a non-graphical solution to Cattell's scree test (Optimal Coordinate criterion) with the nFactors R-package (44).

*Exploratory factor analysis (EFA)*

After identifying the number of shared latent factors in PCA, we performed exploratory factor analysis (EFA), using lavaan, to inform factor loading restrictions and starting values for CFA. We pursued two EFA approaches to estimate factor structures, allowing for the presence of factor correlations or not, using oblique (oblimin) and orthogonal (varimax) factor rotations, respectively. If factor correlations using oblimin were modest (i.e. $r \leq 0.32$)(45), we selected varimax-based rotation to carry out EFA. Factor loadings were selected based on the recommended selection threshold of $|\lambda|<0.1$ (45).

For genetic and residual EFA, we studied the Cholesky-derived genetic and residual covariance, respectively. To account for variation in EFA estimates, given that genetic and residual covariance are estimated with error, we estimated EFA factor solutions using the Diagonally Weighted Least Squares (DWLS) algorithm. This method allows weighting the estimated likelihood function such that any parameter θ is given as

$$\boldsymbol{l}(\boldsymbol{\theta}) = \frac{\mathbf{1}}{\mathbf{2}} \boldsymbol{tr}\left[\left(\boldsymbol{S} - \boldsymbol{\Sigma}(\boldsymbol{\theta})\right)\mathbf{W}^{-\mathbf{1}}\right] \quad [1]$$

where $S$ is the Cholesky-predicted covariance matrix and $\Sigma$ the EFA model-implied genetic covariance matrix. Inverse weighting was carried out with a diagonal weight matrix W, based on the estimated variance $\tilde{V}$ of the genetic covariance VA, as derived with a Cholesky model, where W = diag($\tilde{V}$(VA)).

*Multivariate phenotypic SEM*

We performed phenotypic SEM analogous to the genetic and residual modelling pipeline, as described in the Results. Accordingly, the number of latent phenotypic dimensions was determined with PCA based on the observed phenotypic Pearson's correlation matrix (r:base, pairwise-complete observations of transformed phenotypes). Next, EFA was performed to define CFA model parameters (r:lavaan, v.0.6-11, ML estimator). If shared latent factor correlations (as estimated with oblimin-rotations) were modest ($r \leq 0.3$), we carried out EFA based on varimax-rotations assuming uncorrelated factors. Factor variances were standardised, and factor loadings were restricted to zero if factor loadings were negligible ($|\lambda_{EFA}|<0.1$ (45). EFA and CFA were performed using Pearson's covariance matrices (r:base, pairwise-complete observations of transformed phenotypes) based on split samples, each balanced for missingness. Each model fit was assessed using comparative LL, AIC, and BIC, incremental CFI and TLI (≥0.95) and absolute SRMR (≤0.08) and RMSEA (≤0.06) indices (46).

*GRM-SEM bifactor models*

Bifactor models were fitted to test the independence of identified genetic and residual factors. Benefiting from rotational invariance and unlimited dimensionality, a bifactor model defines one general factor loading on all traits, while any additional factor defines the grouping of traits (47,48). IPC-bifactor compositions were used to assess genetic factorial independence. As for the defined IPC model, IPC-bifactor compositions were informed by consecutive PCA and EFA analyses with the estimated genetic correlation and covariance matrices, respectively. The number of fitted IPC-bifactor models corresponded with the number of latent genetic factors such that IPC-bifactor model 1 refers to the IPC-bifactor model where the first latent factor defines a general factor loading on all traits while any other additional shared latent factor loadings are restricted as determined during EFA-varimax analyses (based on $|\lambda_{EFA}|<0.1$ (45)). IPC-bifactor model 2, in turn, refers to the IPC-bifactor model where the second latent factor defines a general factor loading, etc. IP-/CIP-bifactor compositions were used to assess residual factorial independence. Here, the number of fitted IP-/CIP-bifactor compositions corresponded with the number of latent residual factors as determined during consecutive PCA and EFA analyses with the estimated residual correlation and covariance matrices, respectively. As described for the IPC-bifactor model above, IP-/CIP-bifactor model numbering indicated which latent residual factor defined a general factor loading on all traits. For IP-bifactor models, the corresponding loading restrictions of the latent genetic factors were defined based on the best-fitting IPC model. For the CIP-bifactor models, latent genetic factors formed a Cholesky decomposition.

*<u>GRM-SEM standardized root mean square residual (SRMR)</u>*

As a measure of absolute fit (i.e. independent of log-likelihood and degrees of freedom), we assessed the Standardized root mean square residual (SRMR) (46)

$$\boldsymbol{SRMR}.\mathbf{grm}\boldsymbol{sem} = \sqrt{\frac{\sum_{i=i}^{k}\sum_{j=1}^{i}\left[\frac{s_{ij}-\hat{\sigma}_{ij}}{s_{ii}s_{jj}}\right]^2}{k\,(k+1)}} \quad [2]$$

such that the squared SRMR represents the scaled and squared differences between observed ($s_{ij}$) and estimated ($\hat{\sigma}_{ij}$)trait covariances averaged across the number of free parameters, where the number of free parameters in GRM-SEM is defined as $(k\ (k+1))$. $s_{ii}$ and $s_{jj}$ are the observed standard deviations for trait i and j, respectively, and k represents the number of observed traits. SRMR values >0.08 were considered a bad fit (46).

*<u>Assessment of GRM-SEM genetic or residual trait variance</u>*

We assessed the proportion of phenotypic variance attributable to either genetic (SNP-$h^2$) or residual influences ($e^2$). The total genetic and residual variance of a trait k ($\sigma^2_{g_k}$ and $\sigma^2_{e_k}$, respectively) is given by the sum of the squared standardised loadings of the $j^{th}$ genetic or residual factor (as defined in $\Lambda_A$ and $\Lambda_E$), such that $\sigma^2_{g_k} = \sum_{j=1}^{n} \lambda^2_{a_{kj}}$ and $\sigma^2_{e_k} = \sum_{j=1}^{n} \lambda^2_{e_{kj}}$. Restricted, and thus not modelled, factor loadings were set to null.

*<u>Assessment of proportion of GRM-SEM genetic or residual trait variance explained by latent factors</u>*

We derived the proportion of genetic or residual trait variance that can be explained by a latent genetic or residual factor as factorial co-heritability ($fc_{SNP\text{-}h^2}$) and factorial co-environmentality ($fc_{e^2}$) as

$$fc_{SNP-h^2} = \frac{\lambda^2_{a_{kj}}}{\sigma^2_{g_k}}$$

and

$$fc_{e^2} = \frac{\lambda^2_{e_{kj}}}{\sigma^2_{e_k}},$$

Respectively. Standard errors (SEs) were derived using the Delta method, and *P*-values approximated based on a Wald test.

*<u>GRM-SEM genetic and residual trait correlations</u>*

Genetic and residual trait correlations ($r_g$ and $r_e$, respectively) reflect the extent to which two phenotypes share genetic or residual influences at a scale between -1 and 1. For example, $r_g$ and $r_e$ correlations between phenotypes 1 and 2 can be assessed as follows:

$r_g = \frac{\sigma_{g_{12}}}{\sqrt{\sigma_{g_1}^2 \sigma_{g_2}^2}}$ and $r_e = \frac{\sigma_{e_{12}}}{\sqrt{\sigma_{e_1}^2 \sigma_{e_2}^2}}$

### Supplementary Figures

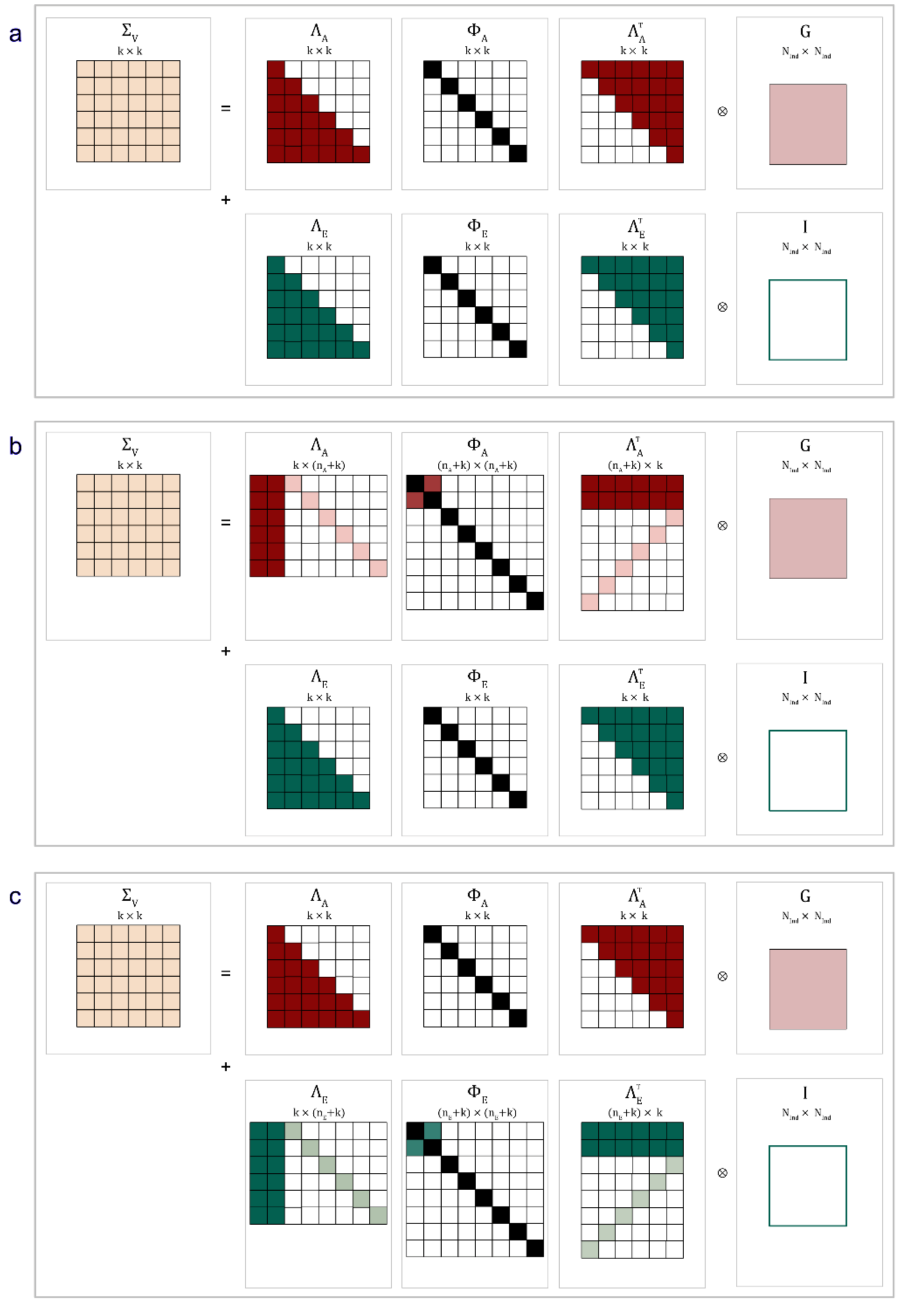

**Supplementary Figure 1**: GRM-SEM Cholesky and hybrid Cholesky models. Schematic representation of GRM-SEM model structures as illustrated for a six-variate trait shown for (a) a Cholesky model, (b) a hybrid IPC model (with 2-factor genetic structure) (c) and a hybrid CIP model (with 2-factor residual structure). $\Lambda_A$ and $\Lambda_E$ capture genetic and residual factor loadings, and $\Phi_A$ and $\Phi_E$ genetic and residual factor (co)variance, with each factor variance constrained to one (i.e. a diagonal of 1). White squares indicate values of zero and black squares values of one.

CIP - Hybrid Cholesky/independent pathway; G - Genetic Relationship Matrix; GRM-SEM - Genetic-Relationship-Matrix Structural Equation Modelling; I – Identity matrix; IPC - Hybrid independent pathway/Cholesky; k - Degrees of freedom; $n_A$ - Number of genetic factor; $n_E$ - Number of residual factors

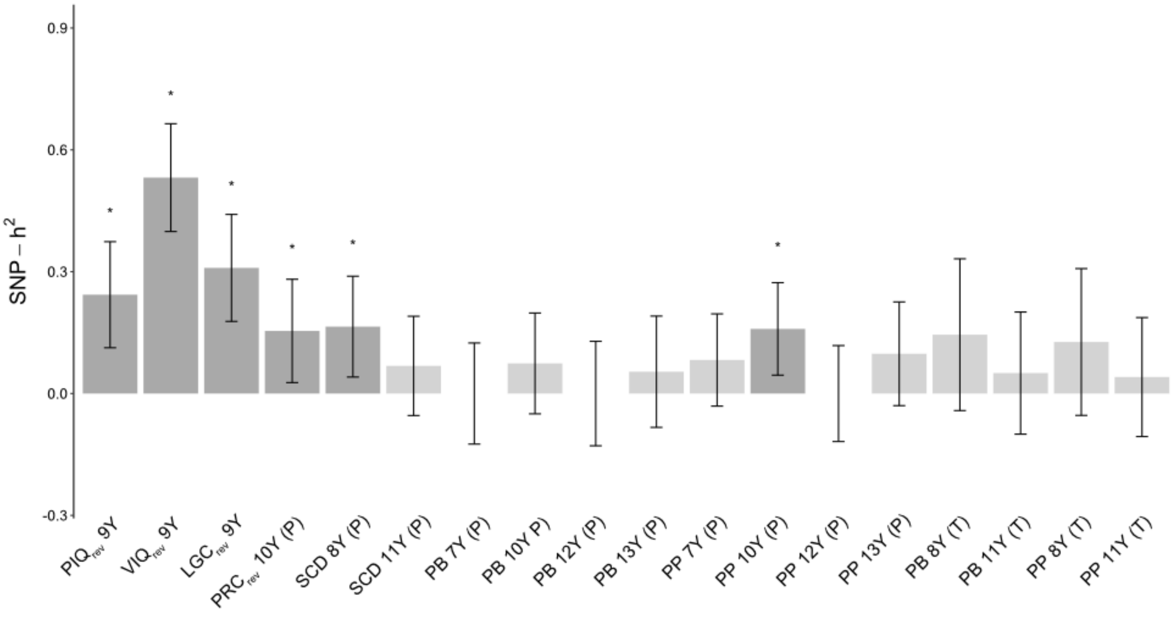

**Supplementary Figure 2: SNP-h²-based phenotype selection in ALSPAC.** Based on a nominal threshold of *P*≤0.05(*), we selected six phenotypes (dark grey) to model the multi-variate structure of cognitive and social/-communication traits during mid-childhood**.** SNP-$h^2$ estimates for transformed measures were estimated with GREML as implemented in GCTA software (v1.26.0).

ALSPAC - Avon longitudinal study of parents and children; LGC - Listening Comprehension; P – Parent-report; PRC - Pragmatic composite scores; PB - Prosocial behaviour; PIQ - Performance intelligence quotient; PP - Peer problems; SCD - Social communication difficulties; SNP-$h^2$ - SNP heritability; T - Teacher-report; VIQ - Verbal intelligence quotient; Y - Years

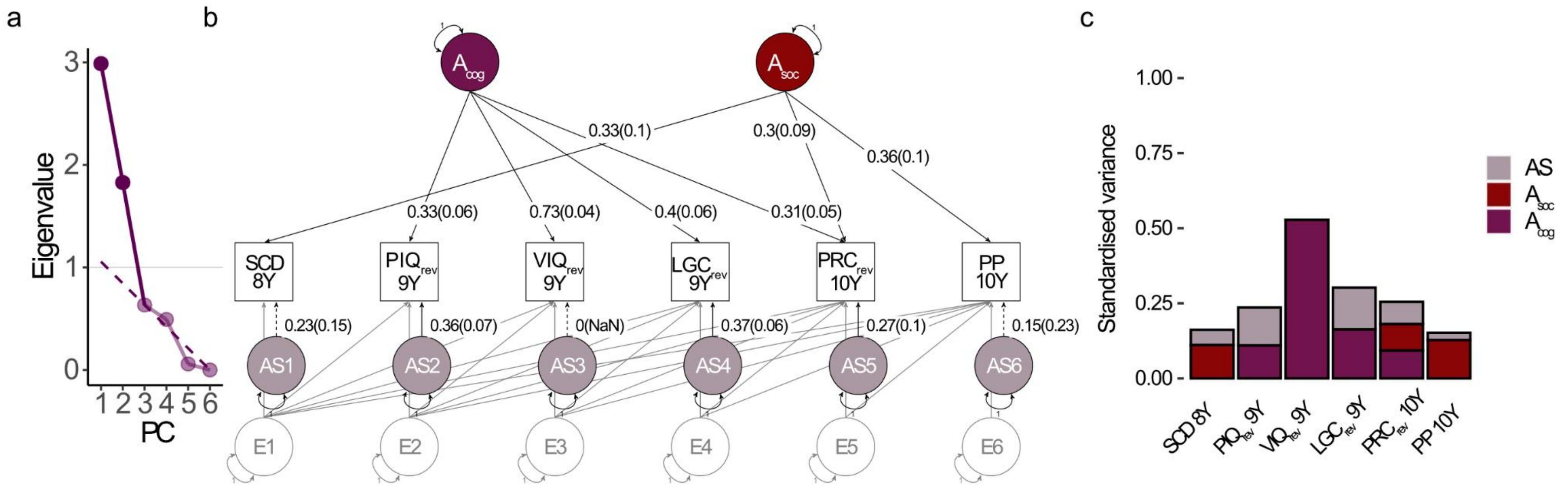

**Supplementary Figure 3: The multi-variate genetic structure between mid-childhood cognitive, language and social traits in ALSPAC. (a)** Two genetic factors were predicted with PCA (a) using Kaiser's criterion (Eigenvalue>1; grey line), and Cattell's scree test (dashed line). Based on an IPC-model, the multi-variate genetic covariance structure was dissected into genetic factors (A) and specific genetic factors (AS) while describing the residual covariance structure (E) with a (saturated) Cholesky decomposition model as depicted in a path-diagram in (b), and as standardised variance decomposition in (c).

The IPC- model was defined based on consecutive PCAs and EFAs based on the genetic correlation matrix (PCA) and the genetic covariance matrix (EFA), as estimated using a saturated Cholesky decomposition. Given that EFA using an oblimin-rotation estimated $r \leq 0.3$, AC loading restrictions were determined based on EFA using a varimax-rotation (restricting ACs to be uncorrelated). The factor loading selection threshold was $\lambda \geq 0.1$.

PIQ, VIQ, LGC, and PRC were reverse-coded (rev) such that a higher score always indicated lower performance/increased difficulties.

ALSPAC - Avon Longitudinal study of Parents and Children; EFA - Exploratory factor analysis; IPC - Hybrid independent pathway/Cholesky; LGC - Listening Comprehension (Wechsler Objective Language Dimension); PRC - Pragmatic composite scores (Children's Communication Checklist); PC - Principal component; PCA - Principal component analysis; PIQ - Performance intelligence quotient (Wechsler Intelligence Scale for Children III); PP - Peer problems (Strengths-and-Difficulties questionnaire); SCD - Social Communication difficulties (Social Communication Disorders Checklist; VIQ - Verbal intelligence quotient (Wechsler Intelligence Scale for Children III); Y - Years

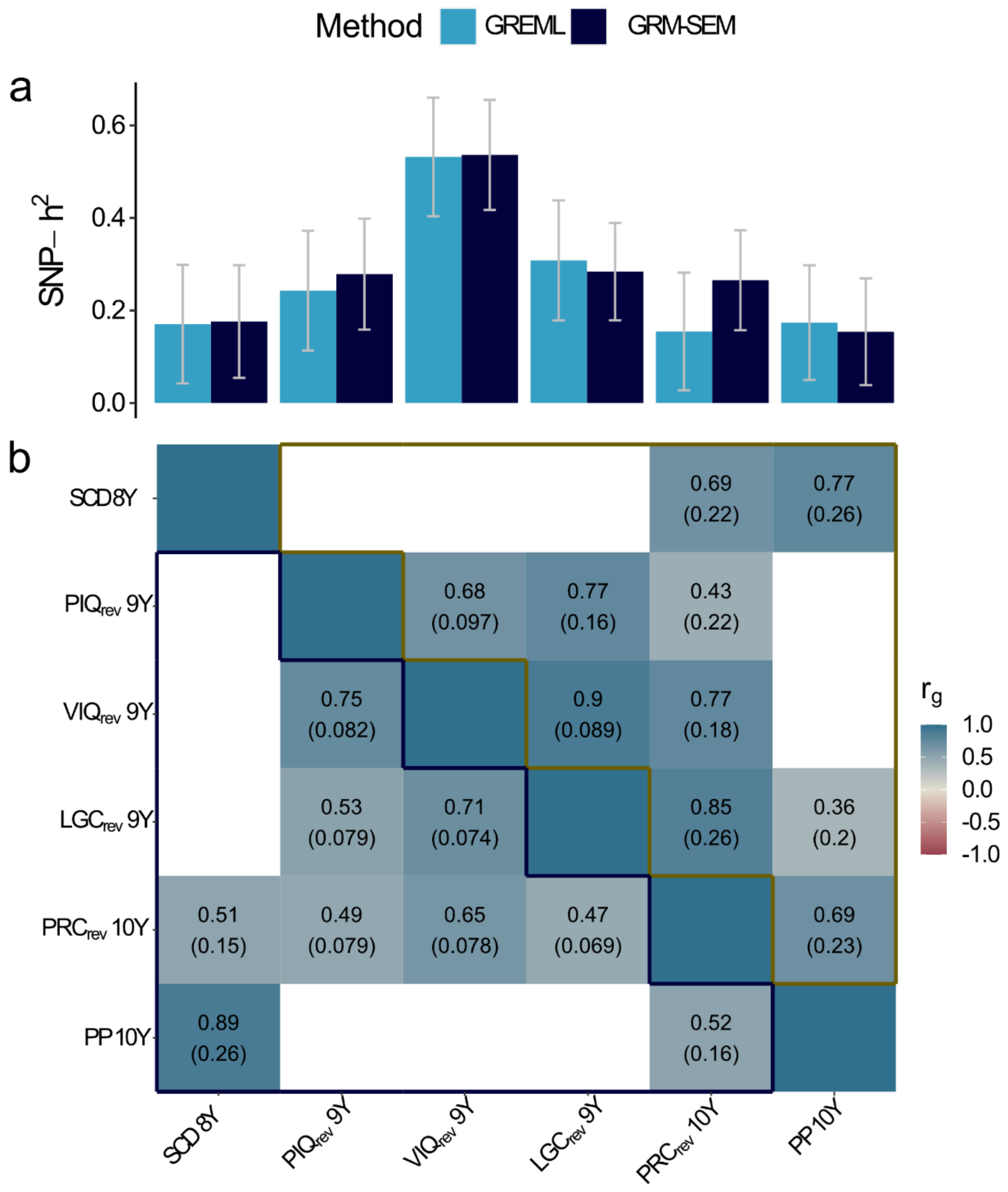

**Supplementary Figure 4: Comparison between GRM-SEM- and GREML-based SNP-h² (a) and $r_g$ (b) estimates of cognitive, language and social traits during mid-childhood in ALSPAC.**

The best-fitting IP model (without $r_{GE}$) was used to derive GRM-SEM-based estimates. Presented GREML-based estimates passed the nominal selection threshold of $P \leq 0.05$. GREML analyses were performed with GCTA software.

PIQ, VIQ, LC, and PAC were reverse-coded (rev) such that a higher score always indicated lower performance/increased difficulties. SCDC, PAC, and PP were based on parent-reports, while PIQ, VIQ, and LC were assessed by a trained interviewer.

ALSPAC - Avon longitudinal study of parents and children; GCTA - Genome-wide complex trait analysis; GREML - Genomic Restricted Maximum likelihood; IP - Independent pathway; LGC - Listening comprehension (Wechsler objective language dimension); PRC - Pragmatic composite (Children's communication checklist); PIQ - Performance intelligence quotient (Wechsler intelligence scale for children III); PP - Peer problems (Strengths-and-difficulties questionnaire); $r_g$ - Genetic correlation; SCD - Social communication difficulties (Social Communication Disorder Checklist); SNP-h² - SNP heritability; VIQ - Verbal intelligence quotient (Wechsler intelligence scale for children III); Y - Years

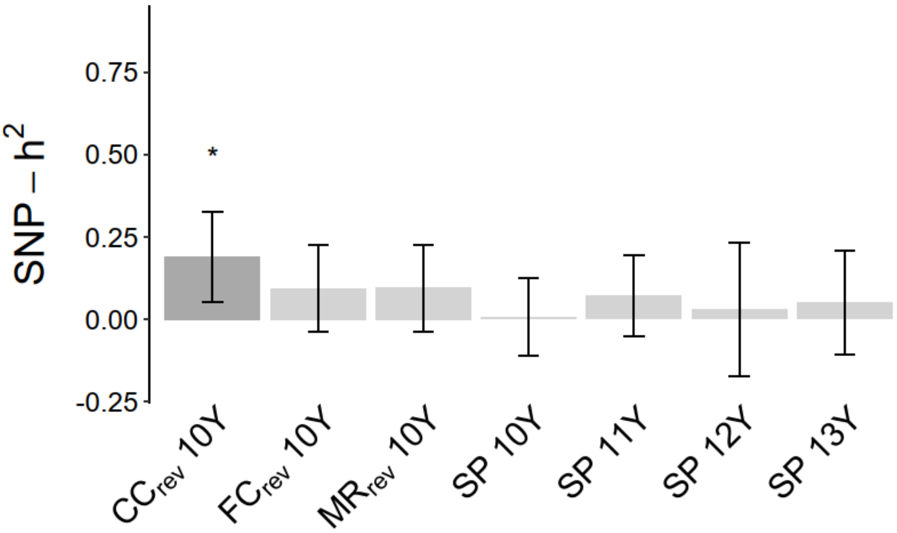

**Supplementary Figure 5: SNP-h²-based phenotype screening of cognitive and social/-communication traits in ABCD.** Phenotypes with nominal evidence for SNP-h² (*P*≤0.05; *) are indicated in dark-grey.
SNP-$h^2$ for transformed measures were estimated with GREML using GCTA software (v1.26.0).

ABCD - Adolescent Brain Cognitive Development study; CC - Crystalized cognition (NIH toolbox); FC - Fluid cognition (NIH toolbox); GCTA - Genome-wide complex trait analysis; MR - Matrix reasoning (Wechsler intelligence test for children V); SNP-h² - SNP heritability; SP - Social problems (Child behaviour checklist); Y - Years

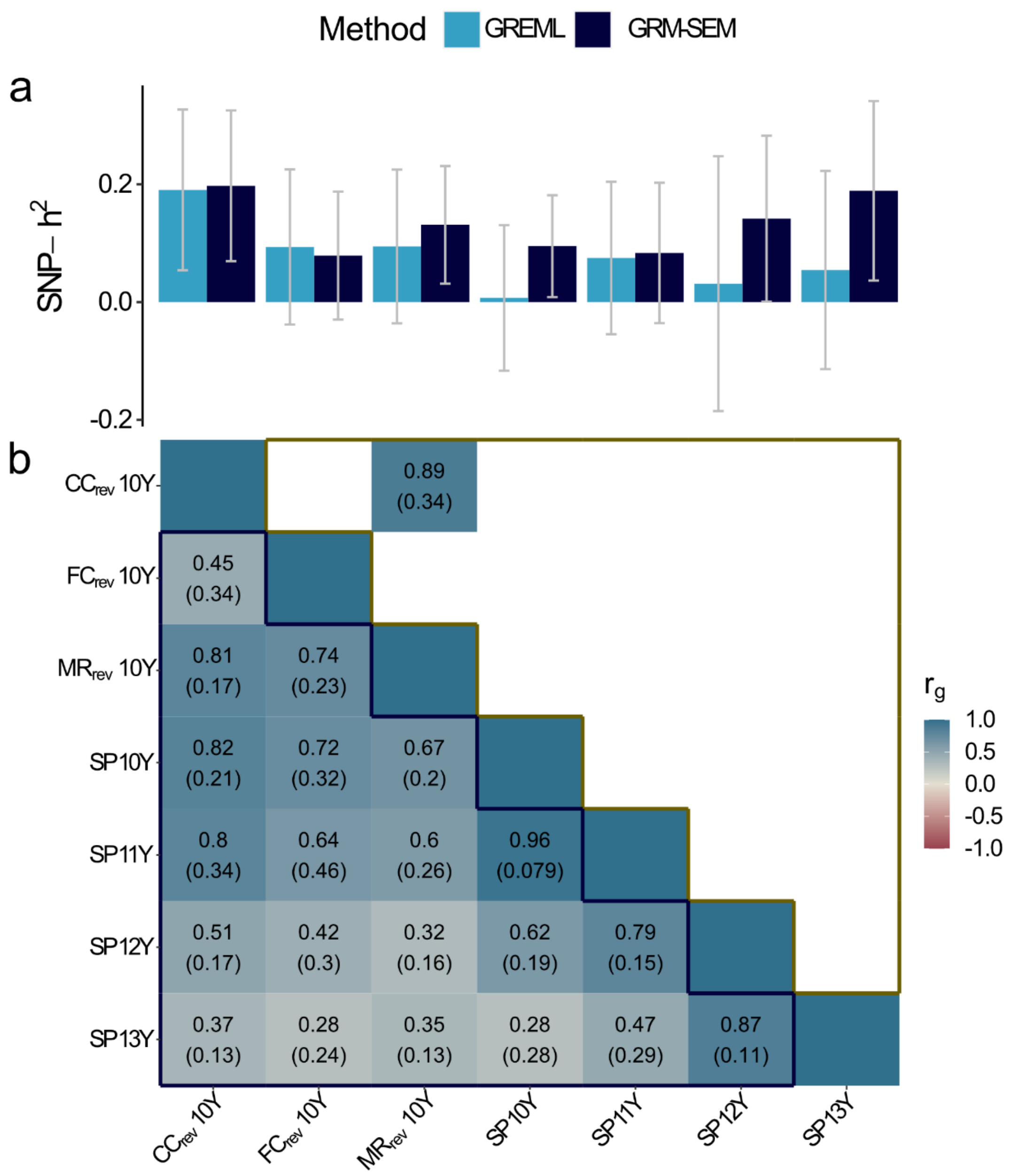

**Supplementary Figure 6: Comparison between GRM-SEM- and GREML-based SNP-h² (a) and $r_g$ (b) estimates of cognitive, language and social traits during mid-childhood in ABCD.**

The best-fitting CIP model was used to derive GRM-SEM-based estimates. Presented GREML-based estimates passed the nominal selection threshold of $P \leq 0.05$. GREML analyses were performed with GCTA.CC, FC, and MR were reverse-coded (rev) such that a higher score always indicated lower performance/increased difficulties. SP were based on parent-reports, while CC, FC, and MR were assessed by a trained interviewer.

ABCD - Adolescent Brain Cognitive Development study; CC - Crystalized cognition (NIH Toolbox); CIP - Hybrid Cholesky/Independent pathway; FC - Fluid cognition (NIH Toolbox); GCTA - Genome-wide Complex Trait Analysis; GREML - Genomic Restricted Maximum likelihood; MR - Matrix reasoning (Wechsler Intelligence Test for Children-V); $r_g$ - Genetic correlation; SNP-h² - SNP heritability; SP - Social problems (Child Behaviour Check List); Y - Years
